# Sleep waves in a large-scale corticothalamic model constrained by activities intrinsic to neocortical networks and single thalamic neurons

**DOI:** 10.1101/2022.10.31.514504

**Authors:** Martynas Dervinis, Vincenzo Crunelli

**Affiliations:** Neuroscience Division, School of Bioscience, Cardiff University, Museum Avenue, Cardiff CF10 3AX, UK

**Author notes:** School of Physiology, Pharmacology and Neuroscience, Biomedical Building, Bristol BS8 1TD, UK. **Authors’ statement** The model codes are available from the corresponding authors upon request.

**Keywords:** sleep spindles, delta waves, slow (<1 Hz) waves, biophysical model, thalamic reticular nucleus

## Abstract

**Aim:** Many biophysical and non-biophysical models have been able to reproduce the corticothalamic activities underlying different EEG sleep rhythms but none of them included the known ability of neocortical networks and single thalamic neurons to generate some of these waves intrinsically.

**Methods:** We built a large-scale corticothalamic model with a high fidelity in anatomical connectivity consisting of a single cortical column and first- and higher-order thalamic nuclei. The model is constrained by different neocortical excitatory and inhibitory neuronal populations eliciting slow (<1 Hz) oscillations and by thalamic neurons generating sleep waves when isolated from the neocortex.

**Results:** Our model faithfully reproduces all EEG sleep waves and the transition from a desynchronized EEG to spindles, slow (<1 Hz) oscillations and delta waves by progressively increasing neuronal membrane hyperpolarization as it occurs in the intact brain. Moreover, our model shows that slow (<1 Hz) waves most often start in a small assembly of thalamocortical neurons though occasionally they originate in cortical layer 5. Moreover, thalamocortical neuron input increases the frequency of slow (<1 Hz) waves compared to those generated by isolated cortical networks.

**Conclusion:** Our simulations challenge current mechanistic understanding of the temporal dynamics of sleep wave generation and suggest testable predictions.

## INTRODUCTION

Recurrent activity in corticothalamic networks is key for the expression of EEG waves of natural sleep.^1-6^ In particular, increases in neuronal membrane hyperpolarization resulting from the vigilance state-dependent, progressively decreasing firing of cholinergic, histaminergic, noradrenergic, and serotoninergic afferents from the hypothalamus and the brain stem underlies the transition from the wake state to sleep spindles, delta and slow (<1 Hz) waves.^7-10^ To further our knowledge of the mechanisms underlying these EEG rhythms, many biophysical and non-biophysical models have been used to simulate sleep waves.^11-21^ However, none of these models implemented the variety of neocortical neuron-specific sleep-related activities reported to occur in the isolated neocortex^22^ and/or the ability of single thalamic neurons to elicit slow (< 1Hz) oscillations when isolated from the neocortex^23-26^.

Using current knowledge of anatomical connectivity^27-38^, we built a large-scale corticothalamic model that contains a single cortical column (with different excitatory and inhibitory neuronal populations) that is reciprocally connected to first-order and higher-order thalamic nuclei and its respective γ-aminobutyric acid (GABA)ergic neurons of the nucleus reticularis thalami (NRT). The model is constrained by implementing the ability of different neocortical populations to generate slow (<1 Hz) oscillations when detached from the thalamus^22,39^ and single thalamic neurons to elicit sleep waves when isolated from the neocortex^23-25^.

Our corticothalamic model accurately reproduces all sleep EEG rhythms and the transition from a desynchronized EEG to various sleep rhythms. Our results shows that the EEG slow (<1 Hz) waves are most often initiated by a small assembly of thalamocortical neurons though they can also originate in cortical layer 5 (L5) and that their frequency is decreased in the absence of thalamic afferents activity. Moreover, our simulations suggest testable predictions.

## METHODS

### Model architecture

The corticothalamic model contains 900 model neurons and is organised into 6 sectors (Fig. S1): four cortical layers, including layers 2/3 (L2/3), 4 (L4), 5 (L5) and 6 (L6), and a first- and a higher-order thalamic nucleus with their thalamocortical neurons (TC_FO_ and TC_HO_, respectively) which are reciprocally connected to inhibitory NRT neurons (NRT_FO_ and NRT_HO_, respectively). Each cortical layer is divided in two subsectors, each with 100 excitatory and 50 inhibitory neurons. Cortical excitatory subsectors contain different number of regular spiking^40^ (RS), intrinsically bursting^40,41^ (IB), early firing^22^ (EF), repetitive intrinsically bursting^22^ (RIB) and network driver (ND) neurons^22^, whereas inhibitory subsectors have fast-spiking (FS) interneurons^42^ (Fig. S1). The full model is a 2-dimensional stack of subsector neuron rows. The neuron position within a subsector was determined pseudo-randomly.

### Model network connectivity

The model network connectivity and the connection weights are provided in Fig. S1 and Tables S1,S2). Connections were organised topographically with sources and targets located in matching regions of their corresponding structures. A neuron did not synapse onto itself and could only form a single synapse on its target neuron. The number of contacts that a source neuron could form in a target structure was defined by the parameter P (a projection radius) (Table S1). Other key connectivity parameters (e.g., connection weight, postsynaptic potential shape, synaptic transmission latency, synaptic receptors) are listed in Table S2 and Supplementary Methods.

We made several simplifying assumptions. In certain instances, we scaled down differences in anatomical projection radii. This was true for intra-cortical connections simplifying network calibration. Moreover, we scaled down the physiological N-methyl D-aspartate (NMDA) component in cortical synapses in order to eliminate a slow periodicity component in paroxysmal simulations which had not been observed experimentally. NRT cells were connected by gap junctions^43^, i.e. the first- and second-degree neighbours formed single 3 GΩ and 4.5 GΩ junctions, respectively.

### Model neurons

TC and NRT neurons were single-compartment Hodgkin-Huxley models whereas cortical neurons were Hodgkin-Huxley models with separate axosomatic and dendritic compartments.

The equations for both thalamic and cortical neuron models are listed in the Supplementary Methods, that also describe the intrinsic and synaptic membrane currents and their dependence on intracellular ion concentration (where applicable) (Tables S3-S7).

### Simulations

All simulations were carried out in NEURON^44^ on a desktop computer or one of the following computing clusters: the Neuroscience Gateway (NSG) Portal for Computational Neuroscience^45^ or the Cardiff University Life Sciences computing cluster.

### Data analysis

Simulation data were analysed and visualised with the help of custom-written Matlab (MathWorks Inc., USA) routines. The scalp EEG signal produced by the simulations was estimated as described by Bédard et al (2004)^46^ (see Supplementary Methods).

## RESULTS

Firstly, we constrained the model by testing the ability of neocortical networks isolated from the thalamus to elicit slow (<1 Hz) oscillations and of single thalamic neurons to produce sleep spindles, delta waves and slow (< 1Hz) oscillations, as reported in many *in vitro* studies.^22-25,39,47,48^

### Slow (<1 Hz) oscillations in the isolated neocortical network

We were able to reproduce the typical firing patterns observed at different levels of membrane polarization in RS and EF neurons, the pyramidal cell type that shows early firing during slow (<1 Hz) oscillations^22^ (Fig. S2), as well as in IB and FS neurons^22^ (Fig. S3). We also developed models of RIB neurons and of the ND pyramidal neurons that “drive” slow (<1 Hz) oscillations in neocortical slices.^22^. The firing patterns of RIB and ND neurons well matched those observed experimentally^22^ (Fig. S4A), including their input-output curves (Fig. S4B).

When all these neocortical excitatory and inhibitory model neurons were synaptically connected in a cortical column (Fig. S1), a pattern of non-rhythmic, low-amplitude activity was observed in the EEG for low K^+^ leak conductance (g_KL_) values (Table S8) in all component neurons (Fig. 1L,M). When g_KL_ was increased (Table S8), however, the neocortical network generated clear EEG slow (<1 Hz) waves (Fig. 1C-E) that could also give rise to faster waves within the delta frequency band (Fig. 1F-H). Thus, all neurons in all cortical layers oscillated at a frequency of 0.44 Hz (Fig. 1C-E), i.e. within the slow (< 1Hz) oscillation frequency, whereas at lower g_KL_ values the oscillation reached the frequency of 1.57 Hz, i.e. within the δ frequency range (Fig. 1F-H): indeed, as g_KL_ was changed the neocortical oscillation frequency smoothly changed accordingly (not shown). Finally, further small progressive decreases of g_KL_ in all cortical cells (except EF neurons) (Table S8) resulted in the break-down of the regular pattern of oscillations (Fig. 1I-K), and eventually led to a desynchronized, low-amplitude EEG similar to that of the wake state (Fig. 1L,M) where single neurons fired randomly.

**Figure 1.**
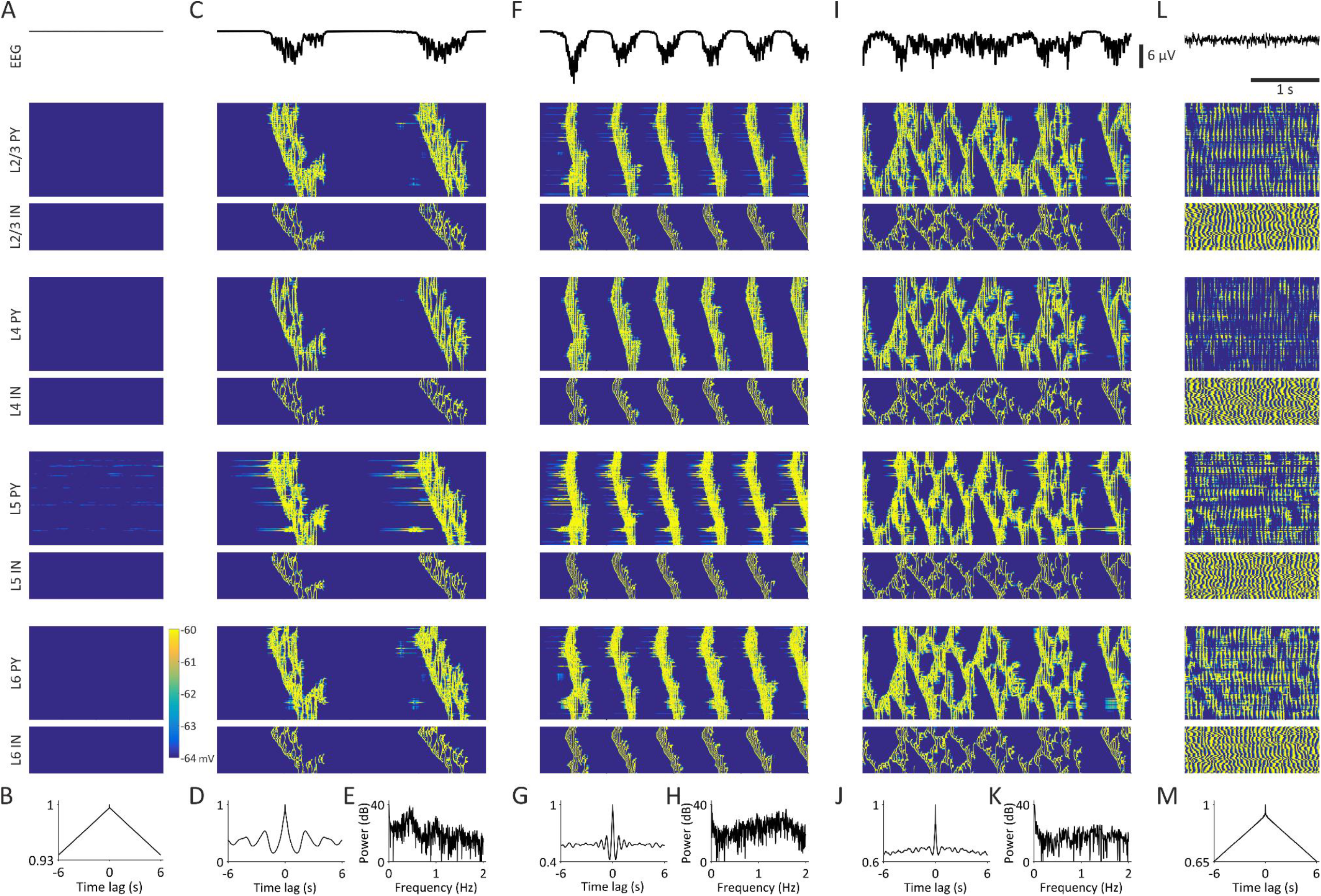
Slow (<1 Hz) and delta oscillations in the isolated neocortical model. EEG (top trace in A,C,F,I,L) and corresponding colour-coded membrane potential graphs for the indicated neuronal populations (bottom traces in A,C,F,I,L), EEG autocorrelograms (B,D,G,J,M) and EEG power graphs (E,H,K) of the indicated activity. A,B, with a large g_KL_ the cortical model does not elicit any activity. C-E, Decreasing g_KL_ of ND and EF cells results in spontaneous slow (< 1Hz) oscillations in the entire cortical network. F-H, Further reducing g_KL_ of ND cells leads to delta waves in the isolated cortical model. I-K, when g_KL_ of other cortical neurons (except ND and EF neurons) is slightly decreased a non-regular activity pattern is observed, i.e. there is a break-down of the oscillations. L,M, The network shows a desynchronized EEG and tonic firing in all neuronal populations when g_KL_ of all cell types (except ND cells) is further decreased. L2/3 PY: pyramidal neurons in cortical layers 2 and 3; L2/3 IN: interneurons in cortical layers 2 and 3; L4 PY: pyramidal neurons in cortical layer 4; L4 IN: interneurons in cortical layer 4; L5 PY: pyramidal neurons in cortical layer 5; L5 IN: interneurons in cortical layer 5; L6 PY: pyramidal neurons in cortical layer 6; L6 IN: interneurons in cortical layer 6.

The slow (<1 Hz) oscillation elicited by the isolated neocortical network (Fig. 1C-E) were initiated in L5 pyramidal neurons since the firing appeared there earlier than in other layers, followed by pyramidal neurons in L6, L2/3 and L4 (Fig. S5A,B). In each layer, the inhibitory neuron firing occurred after that of the pyramidal neurons with little difference among layers (Fig. S5A,B). The ND neurons in L5 are the cells that initiate the oscillation, followed by the EF neurons in L/6 and L2/3 (Fig. S5C). Thus, the onset timing of the simulated neocortical slow (<1 Hz) oscillation is highly consistent with experimental data obtained in cortical slices.^22,39^ Indeed, the duration of the Up- and Down-states of the simulated slow (<1 Hz) oscillations in the isolated neocortex was similar to that observed *in vitro* (∼250 ms and ∼840 ms, respectively)^22^ (Fig. S5D,E).

The two main contributors to the maintenance of the Up-state were, in order of importance, the persistent Na^+^ current (I_Na(P)_) and the excitatory postsynaptic potential (EPSP) barrage in IB (Figs. S6D,F) and RS (not shown) neurons. Reducing α-amino-3-hydroxy-5-methyl-4-isoxazolepropionic acid (AMPA)-mediated EPSPs or I_Na(P)_ abolished Up-states (Fig. S7B,F) confirming their causal roles in the initiation and maintenance of the slow (< 1Hz) oscillation, whereas blockade of NMDA receptors had little impact on Up-states (Fig. S7C).

Other significant contributors in maintaining the Up- and Down-state dynamics were the A-type K^+^ current (I_A_) and the inhibitory postsynaptic potential (IPSP) barrage (Figs. S6F and S7G). Their blockade resulted in over-excitation and the transformation of the slow (<1 Hz) oscillation into a paroxysmal oscillation (Figs. S7D) (implying a balancing role for GABA_A_ receptors) or in the disappearance of the Down-states as the remaining K^+^ currents could no longer terminate Up-states (Fig. S7G) (implying a role for I_A_ in the termination of the slow (<1 Hz) oscillation). The total intrinsic hyperpolarizing and depolarizing currents were similar as were the excitatory and inhibitory synaptic currents (Fig. S6D,E,I).

The termination of the Up-state occurred because of the gradual accumulation of the Ca^2+^-activated K^+^ current (I_K[Ca]_) but also the M-type K^+^ current (I_M_) and the Na^+^-activated K^+^ current (I_K[Na]_) (Fig. S6G). Blocking I_K[Ca]_ in all model neurons except ND increased the Up-state duration confirming its causal role in the termination (Fig. S7I). Similarly, block of I_M_ increased the duration and irregularity of the Up-states confirming its role in curtailing Up-states (Fig. S7J). On the other hand, the block of I_K[Na]_ resulted in an almost-continuous Up-state, i.e. the Up-state never properly terminated (Fig. S7K) since EF neurons fired continuously (not shown).

The contribution to the initiation, maintenance, and termination of Up-states by the T-type Ca^2+^ current (I_T_), the high-threshold Ca^2+^ current (I_HVA_) and the hyperpolarization-activated, cyclic nucleotide-gated current (I_h_) was negligible in IB and RS model cells (Fig. S6H). Blocking I_T_, I_HVA_, and I_h_ (except for ND) had no significant effect on the duration or shape of the Up-states (Fig. S7L-N). However, I_h_ and I_HVA_, as the key pacemaker currents, initiated the Up-states in ND neurons (not shown).

Blocking GABA_A_Rs resulted in a continuous paroxysmal oscillation (Fig. S7D) as it was observed experimentally in cortical slices after application of bicuculline.^49^ Hence, the effect of GABA_A_Rs is to reduce the intensity of cortical Up-states and, in turn, to increase their duration by limiting the build-up of I_K[Ca]_ and I_K[Na]_, an observation consistent with the experimental data.^50^ The role of GABA_B_Rs was to shorten the Up-states since their blockade resulted in Up-states longer than 1 sec (Fig. S6E), an effect opposite to that of GABA_A_R and consistent with *in vitro* experiments.^50^

Having established the dynamics of initiation and maintenance of the EEG slow (<1 Hz) oscillation in the isolated cortical network, we next investigated whether the membrane potential waveforms of the different cortical populations during the Up- and Down-states dynamics were similar to those observed in neocortical slices.^22,39^ The excitatory RS, EF and ND neurons faithfully reproduced the Up- and Down-states dynamics, the firing patterns and the bimodal distribution of the membrane potential characteristic of neocortical slow (<1 Hz) oscillations *in vitro*^22^ (Fig. 2A-H,M-P). In particular, ND neurons exhibited the diverse activity observed in this pyramidal neuron type at an early and late stages of the slow (<1 Hz) oscillation as shown following its pharmacological induction in cortical slices^22^ (Fig. 2O,P). Moreover, the membrane potential waveform of IB and RIB pyramidal neuron models during the simulated neocortical slow (<1 Hz) oscillations well matched that reported in cortical slices^22^ (Fig. S8). A slow (<1 Hz) oscillation was also present in the inhibitory FS neurons, though a clear bimodal distribution was evident in their firing but not in their membrane potential distribution plots as observed experimentally^22^ (Fig. 2I-L). Finally, a faithful reproduction of the slow (<1 Hz) oscillation and its membrane potential distribution plots was also found for IB and RIB neurons^22^ (Fig. S8)

**Figure 2.**
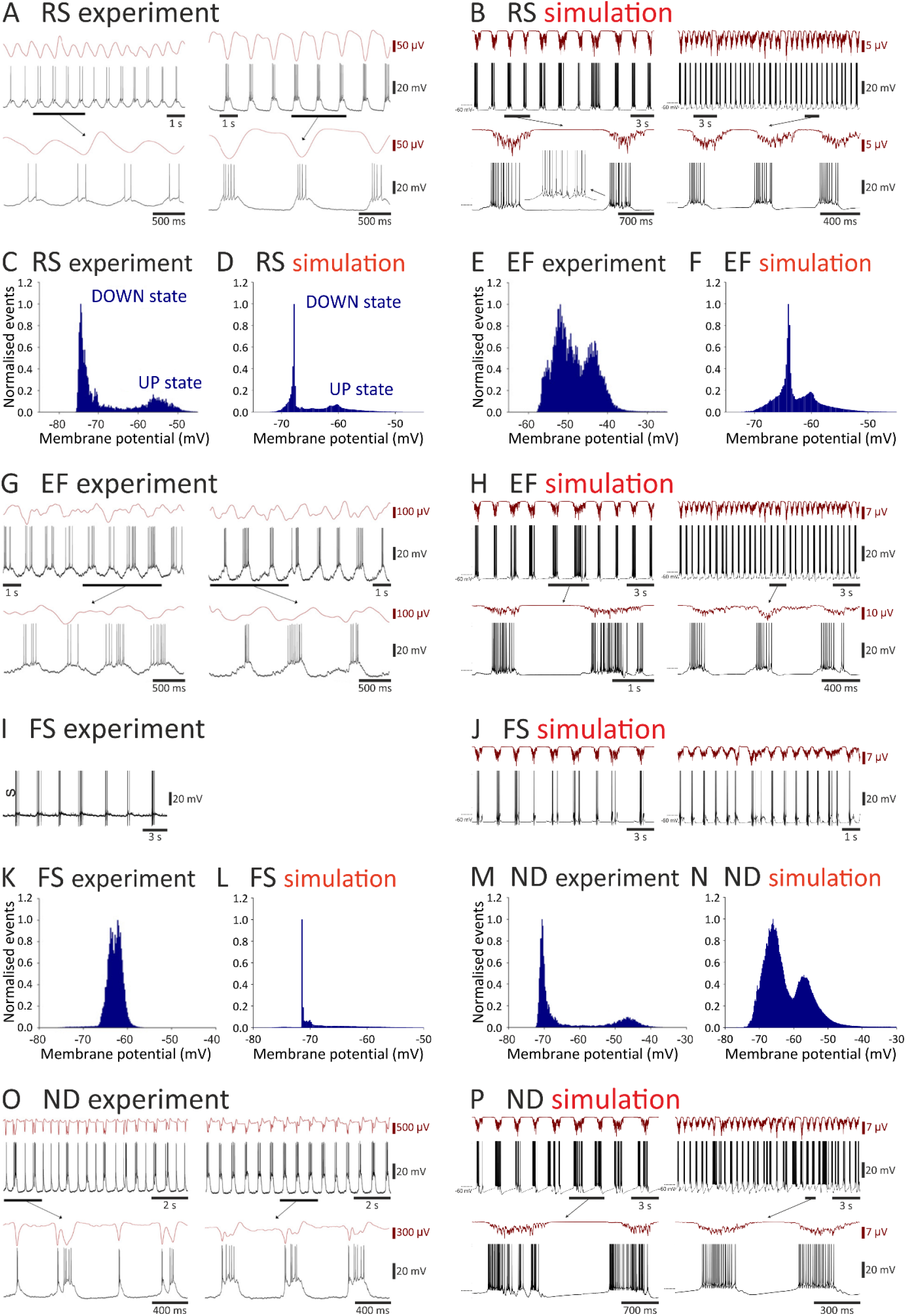
Experimental and simulated membrane potential dynamics in cortical neurons during slow (<1 Hz) oscillations in the isolated cortical network. A,B, Simultaneous local field potential and membrane potential dynamics of an RS neuron recorded *in vitro* (experiment) during slow (<1 Hz) oscillations and its simulated activity in the isolated cortical network (simulation). The left-hand traces show the initial stage of the oscillation while the right-hand traces show the oscillation at a later stage (increased neuromodulatory drive). C,D, Normalized membrane potential distribution plots for the experimental and simulated slow (< 1Hz) oscillations of an RS neuron with the typical peaks of the Up- and Down-states. E,F, Same as C,D but for an EF neuron. G,H, Same as A,B for an EF neuron. I,J, Same as A,B but for an FS neuron. K,L, Same as C,D but for an FS neuron. Note the lack of a clear bimodal membrane potential distribution in the plots of both the experimental and simulated data of the FS neurons. M,N, Same as C,D but for an ND neuron. O,P, Same as A,B but for an ND neuron. Experimental data are reproduced with permission from Lorincz et al. (2015).^22^

### Intrinsic slow (<1 Hz) oscillations of thalamic neurons

Our model TC_FO_, TC_HO_, NRT_FO_ and NRT_HO_ neurons reproduced the ‘classical’ activity patterns of these cells, i.e. tonic single action potential (AP) and T-type Ca^2+^ channel-mediated burst firing, in particular the typical burst signature of decelerando and accelerando-decelerando burst firing patterns of TC and NRT neurons^51^, respectively (not shown).

We then simulated the intrinsic generation of delta and slow (<1 Hz) oscillations observed in TC_FO_ neurons *in vitro* following pharmacological activation of metabotropic glutamate receptors 1a (mGluR1a)^23-25^, which was mimicked by reducing g_KL_ (Fig. 3). The TC_FO_ neurons closely reproduced the hyperpolarization-dependent transition from tonic firing to slow (<1 Hz) oscillations at increasing frequency (due to shorter Up states) and then to delta waves (Fig 3B,D). Moreover, the full spectrum of delta and slow (<1 Hz) oscillations observed at different levels of membrane polarization were elicited by the NRT_FO_ neuron model (Figs. S9). Notably, I_Na(P)_ and the Ca^2+^-activated non-specific cation current (I_CAN_) played a major role in the duration of the Up-state of NRT_FO_ neurons since their reduction led to a progressive shortening, and eventually to the abolition of, the Up-states, i.e. the slow (< 1Hz) oscillation was gradually transformed into a delta wave (Fig. S10B1-C2). Notably, NRT_FO_ neurons were capable of eliciting both slow and fast sleep spindle waves (6.5 - 16Hz) in isolation, i.e. when disconnected from TC_FO_ neurons (Fig. S11). Finally, TC_HO_ and NRT_HO_ neurons reproduced slow (< 1Hz) and delta oscillations similar to those of TC_FO_ and NRT_FO_ neurons, respectively (not shown).

**Figure 3.**
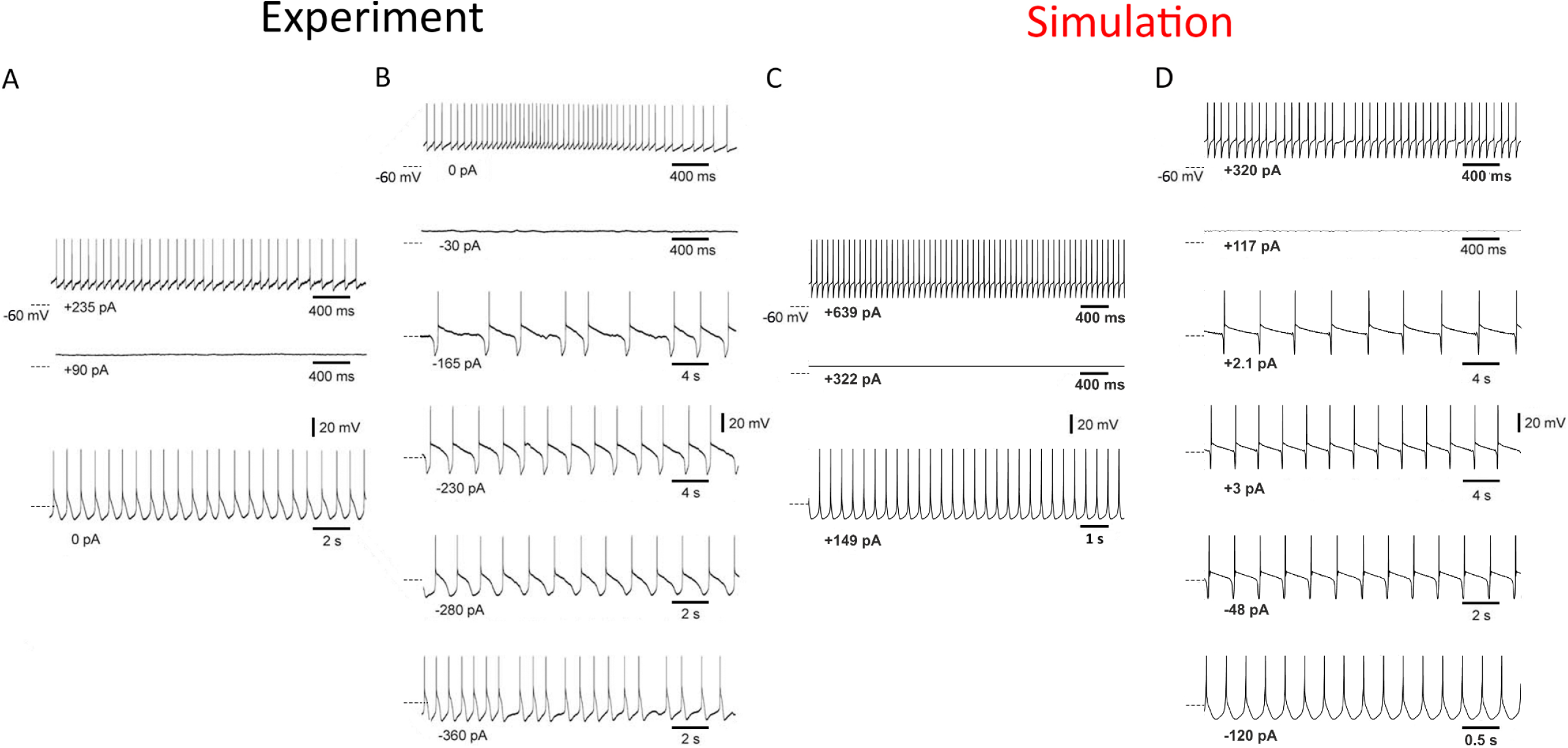
Experimental and simulated membrane potential dynamics of TC_FO_ neurons during intrinsically generated slow (<1 Hz) oscillations. A, Tonic firing (top trace), quiescence (middle trace) and delta oscillations (bottom trace) are observed at different level of membrane polarization (injected current at the bottom of each trace) in a cat ventrobasal TC neuron recorded in a thalamic slice maintained in a standard recording solution. B, After application of 100 μM trans-ACPD (a metabotropic glutamate receptor, mGluR, agonist) the same neuron shows slow (<1 Hz) oscillations between the quiescent state and the delta oscillations (bottom trace). C, Simulation in the TC_FO_ neuron model of the experimental activity shown in A. D, Simulation of experimental activity shown in B was obtained after reducing g_KL_ of the TC_FO_ neuron model to mimic the effect of mGluR activation *in vitro*. Dashed lines on the left of each trace indicate -60 mV. A and B are reproduced with permission from Zhu et al. (2006).^25^

### Sleep waves in the full corticothalamic model

Experimentally it was shown that whereas the neocortex can generate the slow (<1 Hz) oscillation when isolated from the thalamus, its Up-states are less rhythmic and frequent in the absence of the thalamus^52^, a finding that was faithfully reproduced by our full corticothalamic model (Fig. S12). Connecting cortex and thalamus produced stronger Up-state firing in both structures: as a result, K^+^ currents accumulated faster and the Up-states terminated earlier in both structures (not shown). Having shorter thalamic Up-states shortened the whole oscillation cycle because the duration of thalamic Down-states changed little since they are mostly controlled by intrinsic currents. As a result, the next global oscillation cycle was brought forward by an early onset of I_T_-mediated burst firing of TC_FO_ neurons which initiated the Up-states in the cortical neurons (Fig. S12).

Fig. 4A shows simulated slow (<1 Hz) oscillation in the full corticothalamic model with Up-states occurring with a regular periodicity across all neuronal populations. Most commonly (see the first 4 Up-states in Fig. 4B), the Up-state-linked firing appeared initially in a small number of TC_FO_ neurons that, notably, was different from one Up-state to the next (Fig. 4B-C). This was followed by firing in NRT_FO_ and NRT_HO_ neurons and then in L4, L5, L6 and L2/3, with the start of the EEG slow wave coinciding with the start of the L4 neuron firing (Fig. 4C). Notably, when ND cells were depolarised, Up-states could also start in L5 (Fig. S13). TC_HO_ neurons only started firing after all cortical neurons, indicating that cortically driven excitation, more than early inhibition from NRT_HO_ neurons, was necessary to elicit rebound firing at the start of the Up-states in these thalamic neurons (Fig. 4C), as shown experimentally.^23-25^

**Figure 4.**
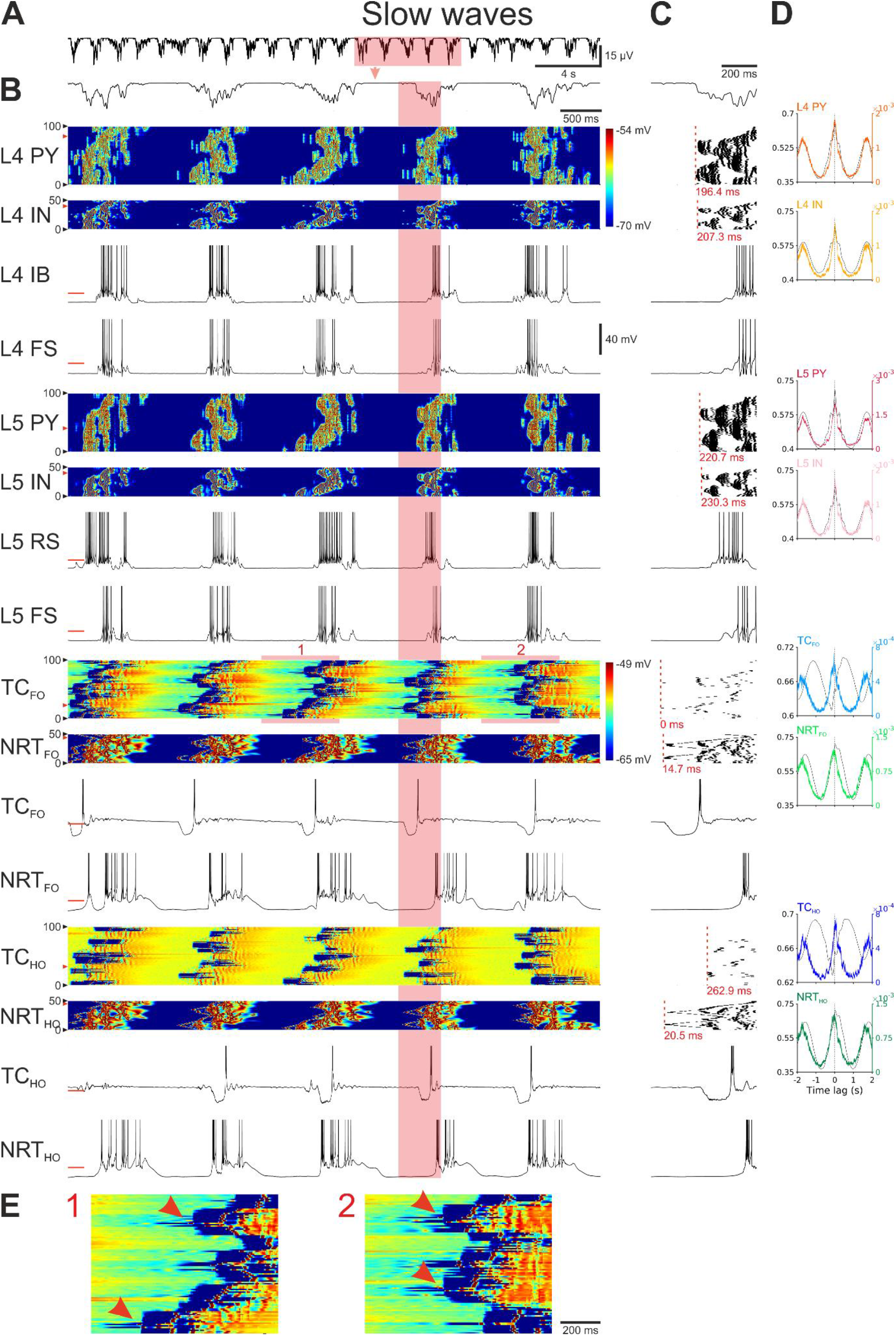
Slow (<1Hz) oscillations in the full corticothalamic model. A, EEG showing the rhythmic pattern of slow (< 1Hz) oscillations. B, EEG (top trace) and colour-coded membrane potential plots of the indicated cortical and thalamic neuronal populations during the 5 cycles of the slow (< 1Hz) oscillation highlighted in A (note the two separate colour-scales for the cortical and thalamic neurons). Below are the corresponding membrane potential waveforms of the two neurons indicated by the red arrow on the left of the corresponding colour-coded plots. C, EEG (top trace), AP rastergrams of the firing in each neuronal population for the slow (< 1Hz) oscillation cycle highlighted in B. Red dashed vertical line represents the first AP of the Up-state in each population. The latency (indicated in red below each rastergram) is measured relatively to the first AP of the cycle in the TC_FO_ neuron that fires first (time zero). Below the rastergrams are the corresponding membrane potential waveforms of that cycle for the indicated neuron. D, Cross-correlations of EEG and membrane potential (black trace) and EEG and APs (colour trace) for the indicated neuronal populations, calculated over a 485 sec-long simulation. Shaded regions are 95% confidence intervals. E, Enlargement of the highlighted sections of the membrane potential colour plot of the TC_FO_ neurons. L4 PY: pyramidal neurons in cortical layer 4; L4 IN: interneurons in cortical layer 4; L4 IB: IB neuron in cortical layer 4; L4 FS: FS neuron in cortical layer 4; L5 PY: pyramidal neurons in cortical layer 5; L5 IN: interneurons in cortical layer 5; L5 RS: RS neuron in cortical layer 5; L5 FS: FS neuron in cortical layer 5; TC_FO_: first order TC neurons; TC_HO_: higher order TC neurons; NRT_FO_: first order NRT neurons; NRT_HO_: higher order NRT neurons.

The membrane potential waveforms of different neuronal types during the slow (< 1Hz) oscillation were similar to those observed experimentally *in vivo*^1,47,53^ (Fig. 4B). The most active Up-states among the cortical neurons were in ND, EF, and FS neurons, whereas RS, IB, RIB displayed on average a sparser firing (Fig. 4B). The firing in thalamic neurons was more regular than in cortex and had a clear intrinsic aspect to it (Fig. 4B). The onset of regularly shaped UP-states in all thalamic neurons followed a Down-state of an almost fixed-length and invariably terminated with an I_T_-mediated burst of action potentials (Figs. 3D and 4A), as observed in thalamic neurons *in vitro*^23-26^: indeed, thalamic activity was essentially an intrinsic oscillation that was shaped by, and synchronized with, the neocortex, a result supported by experimental findings.^26,54^

Notably, when g_KL_ of TC and NRT neurons was increased (Table S9) delta waves started to occur during the Down-states of the slow (<1 Hz) oscillations of these thalamic neurons, as observed experimentally^23-25^: their reflection in the cortical territory resulted in a speeding up of the slow (< 1Hz) oscillation into the delta frequency range (Fig. 5B) as seen in the EEG (Fig. 5A). Whereas during slow (< 1Hz) waves firing was observed first in a small number of TC_FO_ neurons, during delta waves there was little time lag across the entire TC_FO_ neuron population and the delay between thalamic and L4 neuron firing in each cycle was much smaller (<100 msec) (Fig. 5C-F).

**Figure 5.**
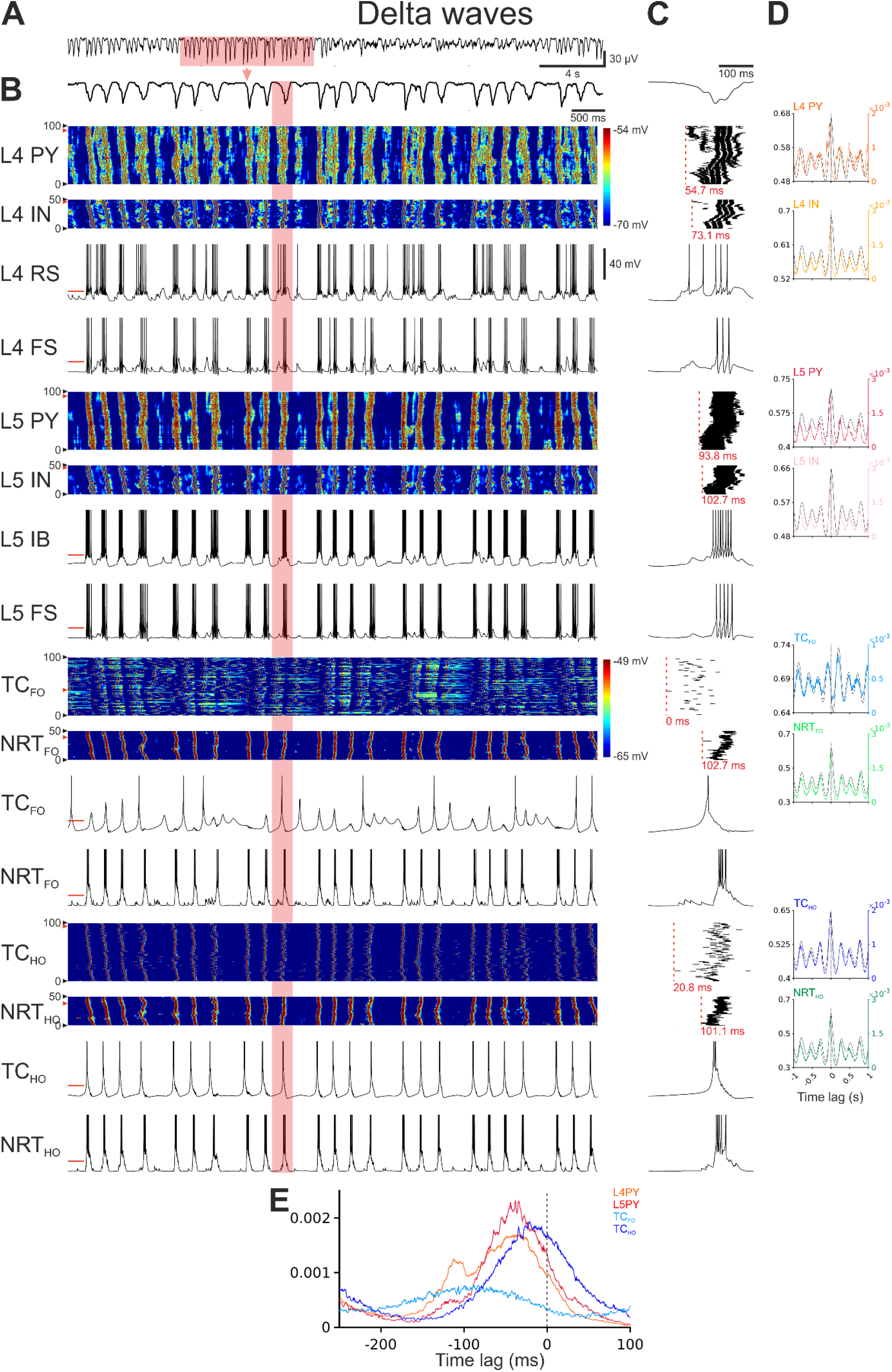
Delta waves in the full corticothalamic model. A, EEG showing a period of rhythmic delta waves. B, EEG (top trace) and colour-coded membrane potential plots of the indicated cortical and thalamic neuronal populations during the section of delta waves highlighted in A (note the two separate colour-scales for the cortical and thalamic neurons). Below are the corresponding membrane potential waveforms of the two neurons indicated by the red arrow on the left of the corresponding colour-coded plots. C, EEG (top trace), AP rastergrams of the firing of each neuronal population for the delta wave cycle highlighted in B. Red dashed vertical line represents the first AP of the Up-state in each population. The latency (indicated in red below each rastergram) is measured relatively to the first AP of the cycle in the TC_FO_ neuron that the fires first (time zero). Below the rastergrams are the corresponding membrane potential waveforms of that cycle for the indicated neuron. D, Cross-correlations of EEG and membrane potential (black trace) and EEG and APs (colour trace) for the indicated neuronal populations, calculated over a 485 sec-long simulation. Shaded regions are 95% confidence intervals. Dashed vertical line indicates zero lag. E, Distribution of the first AP in a delta wave cycle with respect to the EEG for all APs of the indicated neuronal populations. Shaded regions are 95% confidence intervals. Dashed vertical line indicates zero lag. L4 PY: pyramidal neurons in cortical layer 4; L4 IN: interneurons in cortical layer 4; L4 RS: RS neuron in cortical layer 4; L4 FS: FS neuron in cortical layer 4; L5 PY: pyramidal neurons in cortical layer 5; L5 IN: interneurons in cortical layer 5; L5 IB: IB neuron in cortical layer 5; L5 FS: FS neuron in cortical layer 5; TC_FO_: first order TC neurons; TC_HO_: higher order TC neurons; NRT_FO_: first order NRT neurons; NRT_HO_: higher order NRT neurons.

Finally, by adjusting g_KL_ and increasing the fast component of K^+^ afterhyperpolarizing current (I_AHP_) in NRT cells the corticothalamic model was able to reproduce EEG sleep spindles (Figs. 6 and S14). Spindle waves appeared regularly in the EEG and had the “classical” waveform observed *in vivo*^1,47^ (Figs. 6A,B and S14A,B). During a spindle wave, firing started in NRT_FO_ neurons, followed in a few milliseconds by TC_HO_ and NRT_HO_ neurons, and then TC_FO_ and cortical neurons (Fig. 6B-E). Notably, the first firing in TC_FO_ neurons resulted from IPSPs summation leading to a rebound burst, partly explaining the firing initiation in these thalamic neurons (Fig. 6B,E). However, the first firing of a spindle cycle could also originate in TC_HO_ neurons (Fig. S14). Notably, in both cases it is evident that the spindle wave builds up in the thalamus before being reflected into the cortical territory (Figs. 6C and S14B).

**Figure 6.**
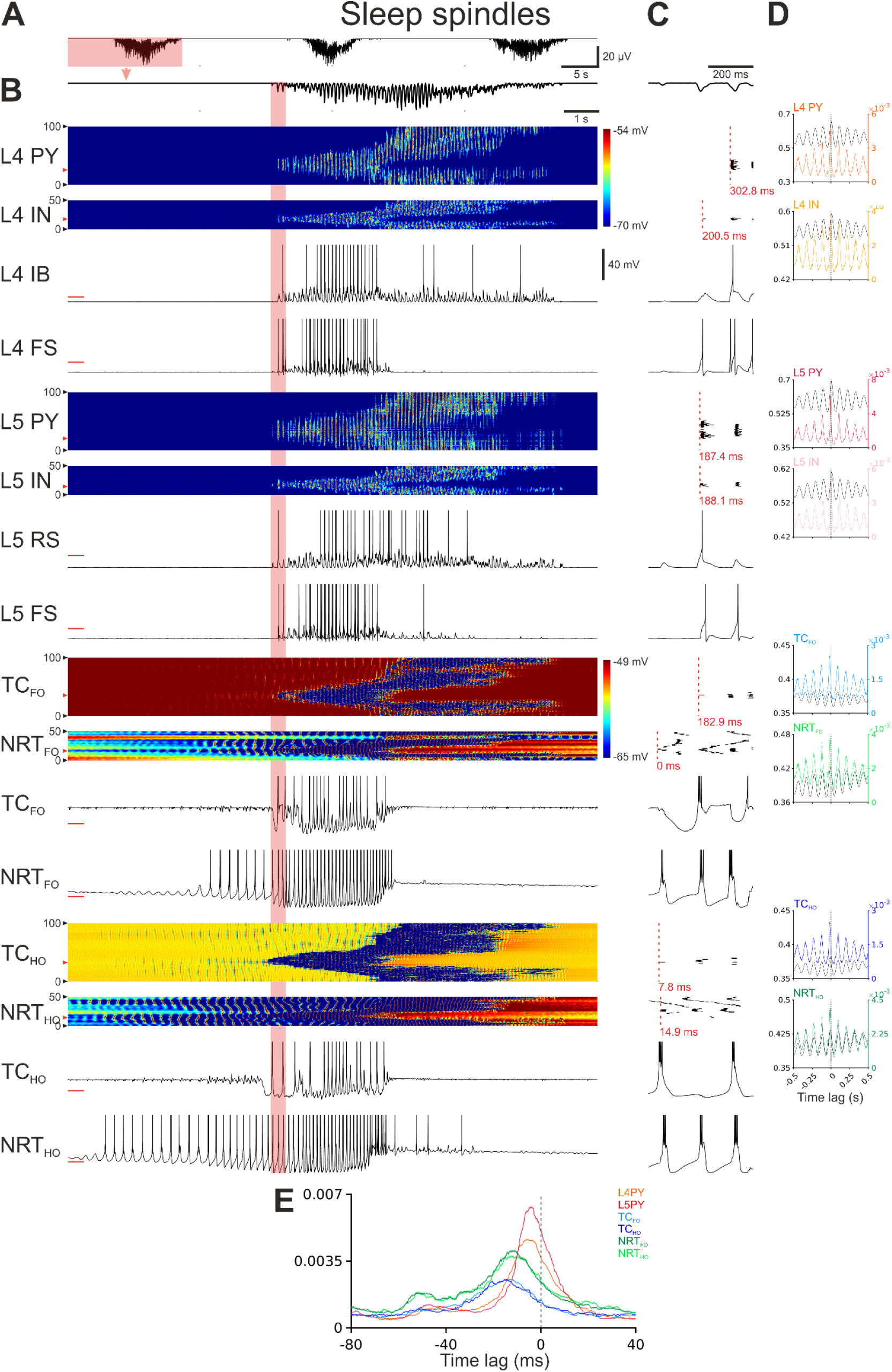
Sleep spindles in the full corticothalamic model. A, EEG showing the rhythmic pattern of sleep spindles. B, EEG (top trace) and colour-coded membrane potential plots of the indicated cortical and thalamic neuron populations during the sleep spindle highlighted in A (note the two separate colour-coded scales for the cortex and the thalamus). Below are the corresponding membrane potential waveforms of the two neurons indicated by the red arrow on the left of the corresponding colour-coded plots. C, EEG (top trace), AP rastergrams of the firing of the first AP in each neuronal population for the onset of sleep spindles highlighted in B. Red dashed vertical line represents the first AP of the Up-state in each population. The latency (indicated below each rastergram) is measured relatively to the first AP of the cycle in an NRT_FO_ neuron (time zero). Below the rastergrams are the corresponding membrane potential waveforms of that cycle for the indicated neuron. D, Cross-correlations of EEG and membrane potential (black trace) and EEG and APs (colour trace) for the indicated neuronal populations, calculated over a 485 sec-long simulation. Shaded regions are 95% confidence intervals. Dashed vertical line indicates the zero lag. E, AP distribution with respect to the EEG for all APs of the indicated neuronal populations. Shaded regions are 95% confidence intervals. Dashed vertical line indicates the zero lag. F, Distribution of the first AP in a spindle cycle with respect to the EEG for all APs of the indicated neuronal populations. Shaded regions are 95% confidence intervals. Dashed vertical line indicates the most negative EEG peak of spindle cycle. L4 PY: pyramidal neurons in cortical layer 4; L4 IN: interneurons in cortical layer 4; L4 IB: IB neuron in cortical layer 4; L4 FS: FS neuron in cortical layer 4; L5 PY: pyramidal neurons in cortical layer 5; L5 IN: interneurons in cortical layer 5; L5 RS: RS neuron in cortical layer 5; L5 FS: FS neuron in cortical layer 5; TC_FO_: first order TC neurons; TC_HO_: higher order TC neurons; NRT_FO_: first order NRT neurons; NRT_HO_: higher order NRT neurons.

## DISCUSSION

The main finding of this study, that used a bottom-up modelling strategy to constrain the corticothalamic model with parameters obtained from *in vivo* and *in vitro* experiments, is the ability to replicate in the EEG and in cortical and thalamic neuronal populations the slow (<1 Hz), delta and spindle waves and the transition between these sleep states by mimicking the neuronal input resistance changes induced by the hypothalamic and brain stem neuromodulatory drives. Indeed, our simulation results remarkably well reproduced the neuronal firing patterns and their sequence of occurrence during different sleep waves, the membrane potential bistability underlying the slow (<1 Hz) oscillation in single cortical and thalamic neurons, the onset and duration dynamics of Up- and Down-states, and the increased regularity and frequency imposed by the thalamus on the slow (<1 Hz) oscillation of the isolated neocortical network.

### Model limitations

The ND neuron model has limitations, including the Down-states being more depolarised and having a larger variance due to a pronounced I_h_-mediated sag than ND neurons recorded *in vitro*.^22^ It may be possible that I_T_ is the main pacemaker in the real ND neurons as opposed to I_HVA_ in the model cells. In the absence of experimental data, initial attempts to model ND neurons used the I_T_ of TC neurons. However, either because I_T_ parameters are not appropriate for the cortical axosomatic and dendritic compartments or the model ND neuron morphology was inadequate, I_T_ did not support the intrinsic slow (<1 Hz) oscillation. Having I_T_ as the actual pacemaker current and not I_HVA_ could drastically stabilize slow (<1 Hz) oscillations by increasing the membrane potential difference between Up- and Down-states in ND neurons and by increasing the AP frequency within a burst leading to more densely-packed EPSPs in the postsynaptic neurons and thus helping the transition to a new Up-state.

The frequency of slow/delta intrinsic oscillations of ND cells increases as they are more depolarised, a result similar to that observed *in vitro*^22^ This, in turn, is reflected in the frequency of modelled cortical network slow/delta oscillations, though this is the opposite of the results in cortical slices.^22^ As indicated above, this is most likely the consequence of the lack of any data on the relative contribution of different intrinsic currents to the excitability of these neurons.

The model does not include the tight regulation of the intrinsic oscillations of thalamic neurons by changes in the function of their metabotropic glutamate receptors as shown experimentally^23-25^, i.e. the ability of these neurons to behave as conditional oscillators.^26^ This could be achieved by linking activation of these modulatory receptors to the firing of the thalamic projecting L5-6 axons, as shown with electrical stimulation of the corticothalamic afferents in thalamic slices.^55^

### Model strengths

Up-states in the cortical network model were initiated by ND and EF neurons, and, to the best of our knowledge, this study is the first that successfully simulate EEG slow (<1 Hz) waves with these intrinsic mechanisms. However, having these intrinsic initiation processes does not exclude the involvement of other mechanisms of initiation and maintenance of this sleep rhythm, in particular the accumulation of spontaneous synaptic activity^56^ and the synchronization of already active neuronal assemblies.^57^ Notably, however, in our model simulated excitatory synaptic conductances were slightly higher than the inhibitory ones, in contrast to experimental studies that reported the opposite^58^ or an equal influence by the two^59^, though these studies were performed *in vitro*.

Thus, our model confirms and enlarges our previous simulations obtained in a thalamocortical model with a limited number of cortical neuron types and fewer thalamic neuron conductances.^19^ Notably, our simulations are also in agreement with the experimental data of David et al. (2013)^52^ as they show that connecting thalamus to cortex increases the frequency and rhythmicity of EEG slow (<1 Hz) waves.

Whereas in the isolated neocortex, the appearance of the slow (< 1Hz) oscillations matches the temporal profile reported in cortical slices^22,39^, i.e. L5, and L4 and L6, in the full corticothalamic model the first *cortical* firing is preferentially observed in L4 followed by L5 and then L6. This is not surprising because of the prevailing input to L4 from the TC_FO_ neurons that are the first to start firing in the full corticothalamic model. However, there are cases in the corticothalamic model where firing starts in L5, as in neocortical slices.^22,39^ Thus, whereas the full corticothalamic model maintains the inherent ability of cortical networks to start slow (< 1 Hz) oscillations in L5, the thalamocortical input most often prevails when the reciprocal cortical and thalamic projection are active.

Finally, connecting thalamus and cortex allows the simultaneous occurrence of different sleep rhythms. In fact, our simulations show that whereas the isolated cortex is capable of generating slow (< 1Hz) and delta oscillations they cannot occur at the same time. In contrast, when the thalamocortical input is intact, slight changes in g_KL_ could smoothly increase the frequency of slow (< 1Hz) oscillations (with a progressive decrease in the UP-state duration), eventually leading to delta waves where as shown experimentally^23-25^ the low threshold calcium potential does not lead to an UP-state, i.e. there is a continuum between slow (< 1Hz) and delta oscillations. Also, sequences of sleep spindles start being expressed cortically and are typically grouped by Up-states. In summary, the richness and versatility of oscillatory behaviours displayed by the full corticothalamic model is, as expected, far greater than any of those generated by its constituent parts, i.e. neocortex and thalamus, in isolation.

### Predictions and future studies

Our simulation results suggest testable predictions that should guide future studies. These may include:

1. temporal dynamics of cortical firing during sleep slow (<1 Hz) waves in non-anaesthetized humans and in naturally waking-sleeping animals;
2. relative contribution of I_Na(P)_ and I_CAN_ to the intrinsic slow (<1 Hz) oscillations of TC and NRT neurons;
3. currents essential to slow (<1 Hz) oscillations in TC and NRT neurons, in particular I_Twindow_, though the large dendritic distribution of I_T^60,61^_ may render this work difficult;
4. re-investigation of the ability of NRT neurons to generate intrinsically sleep spindles in freely moving animals with combined optogenetic and large neuronal assembly recordings and in thalamic slice where the hypothalamic and brain stem drives, known to modulate I_AHP^10^_, are artificially recreated;
5. the pacemaker currents of neocortical neurons, in particular I_HVA_ and I_h_, in ND and EF neurons;
6. relative contribution of I_Na(P)_, I_K[Na]_, and other intrinsic currents to initiation, maintenance, and termination of Up-states in the isolated cortex;
7. increased frequency of corticothalamic Up-states imposed by thalamic drive compared to the isolated cortical network;
8. corticothalamic sleep spindle generation by combination of input resistance and neuromodulators-dependent changes in I_AHP_ of NRT neurons.

## Acknowledgments

This work was supported by an MRC PhD studentship to MD and by the Ester Floridia Neuroscience Research Foundation (grant 1502 to VC).

## Notes

**Conflict of interest** The authors declare no conflict of interest.

### Competing Interest Statement

The authors have declared no competing interest.

