## Supplementary Information for "Sleep waves in a large-scale corticothalamic model constrained by activities intrinsic to neocortical networks and single thalamic neurons"

| | |
|---|---|
| Supplementary Figures | page 2 |
| Supplementary Tables | page 22 |
| Supplementary Methods | page 29 |
| Supplementary Appendices | page 33 |
| Supplementary References | page 47 |

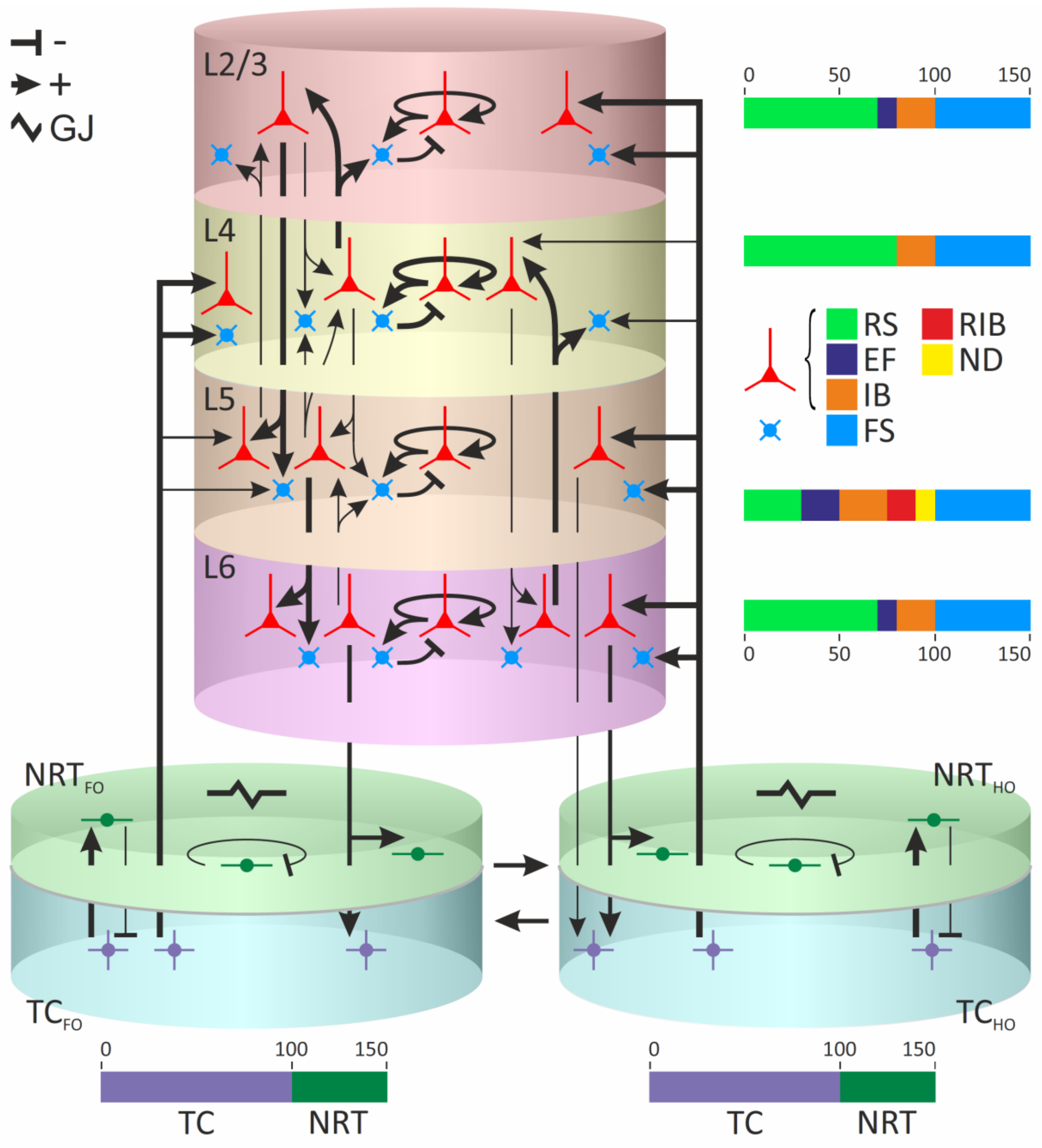


**Figure S1. Corticothalamic model architecture.**

The corticothalamic model consists of 900 neurons distributed in distinctly coloured layers of a single cortical column and two coloured-coded thalamic sectors, a first- and a higher-order sector. Each cortical layer contains 100 excitatory neurons (detailed neuronal populations are in the colour-coded bar on the right) and 50 fast-spiking (FS) inhibitory neurons. Thalamic cylinders (of both first-order and higher-order sector) contain 100 thalamocortical (TC) neurons and 50 nucleus reticularis thalami (NRT) neurons. Sharp and blunt arrows represent excitatory and inhibitory synaptic connections, respectively, with the line thickness indicating the synaptic connection strength (see actual values in Table S1). The lightning symbol indicates gap junctions (GJ) that are present only between NRT neurons.

**L2/3**: cortical layer 2/3

**L4**: cortical layer 4

**L5**: cortical layer 5

**L6**: cortical layer 6

**RS**: regular spiking neuron

**EF**: early firing neuron

**IB**: intrinsically bursting neuron

**RIB**: repetitive intrinsically bursting neuron

**ND**: network driver neuron

**FS**: fast spiking neuron

**$TC_{FO}$**: thalamocortical neurons of first-order thalamic nucleus

**$TC_{HO}$**: thalamocortical neurons of higher-order thalamic nucleus

**$NRT_{FO}$**: nucleus reticularis thalami neurons connected to first-order nucleus,

**$NRT_{HO}$**: nucleus reticularis thalami neurons connected to higher-order nucleus.

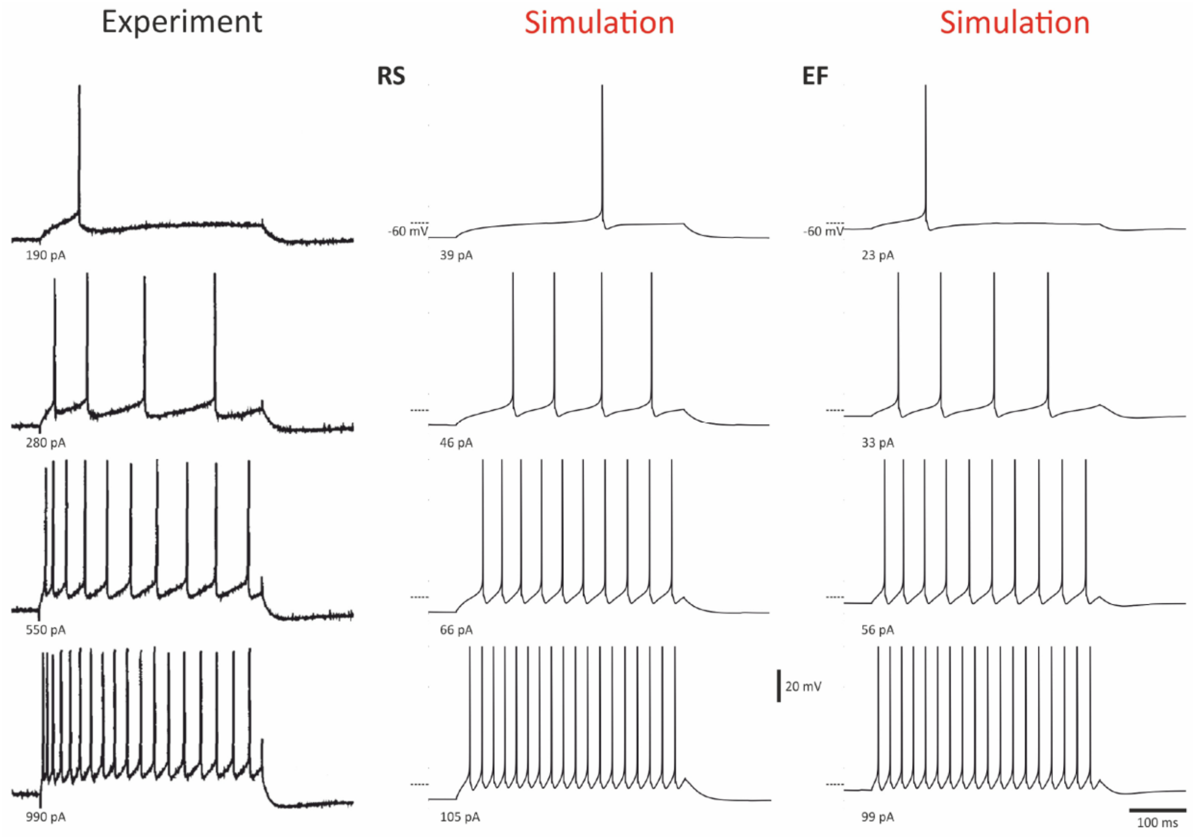


**Figure S2. Experimental and simulated firing patterns of RS and EF neurons.**

Intracellularly recorded firing patterns of a cat motor cortex RS neuron (Experiment, left column), simulated activity of the RS neuron model (Simulation, middle column) and simulated firing of an EF neuron (right column). Value of injected current is indicated below each trace. Note the more depolarized membrane potential of the EF neuron model compared to the RS neuron model, that was achieved by reducing $g_{KL}$ in normal RS model neurons. Dashed line on the left of the traces indicates -60 mV. Experimental data are reproduced with permission from Chen, Zhang, Hu and Wu (1996)[1].

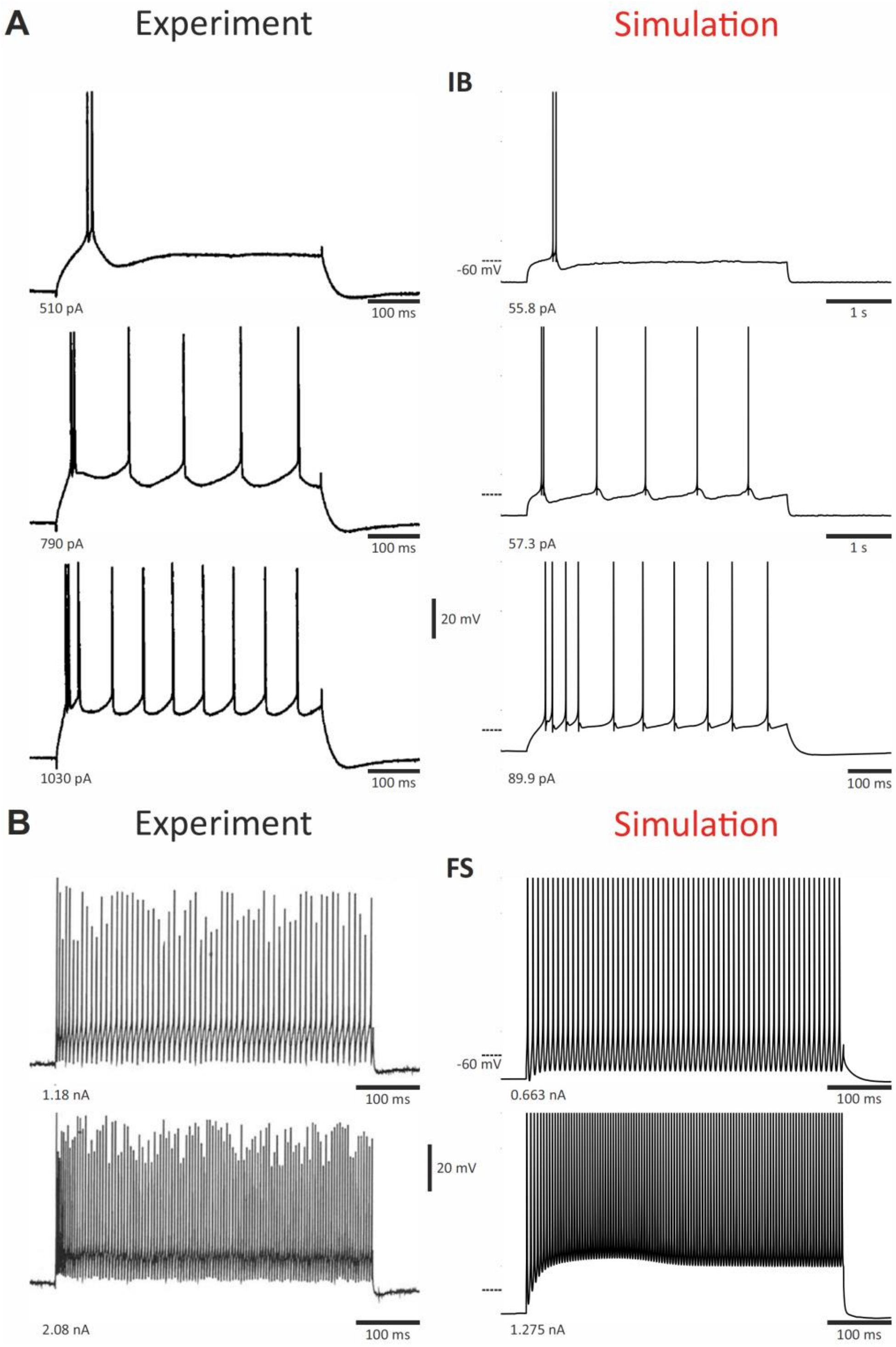


**Figure S3. Experimental and simulated firing patterns of IB and FS neurons.**

A, Intracellularly recorded firing patterns of a cortical RS neuron (Experiment) and simulated activity of the RS neuron model (Simulation). B, Intracellularly recorded firing patterns of a rat cortical FS neuron (Experiment) and simulated activity of the FS neuron model (Simulation). Value of injected current is indicated below each trace. Dashed line on the left of the traces indicates -60 mV. Experimental data are reproduced with permission from Chen, Zhang, Hu and Wu (1996)[1].

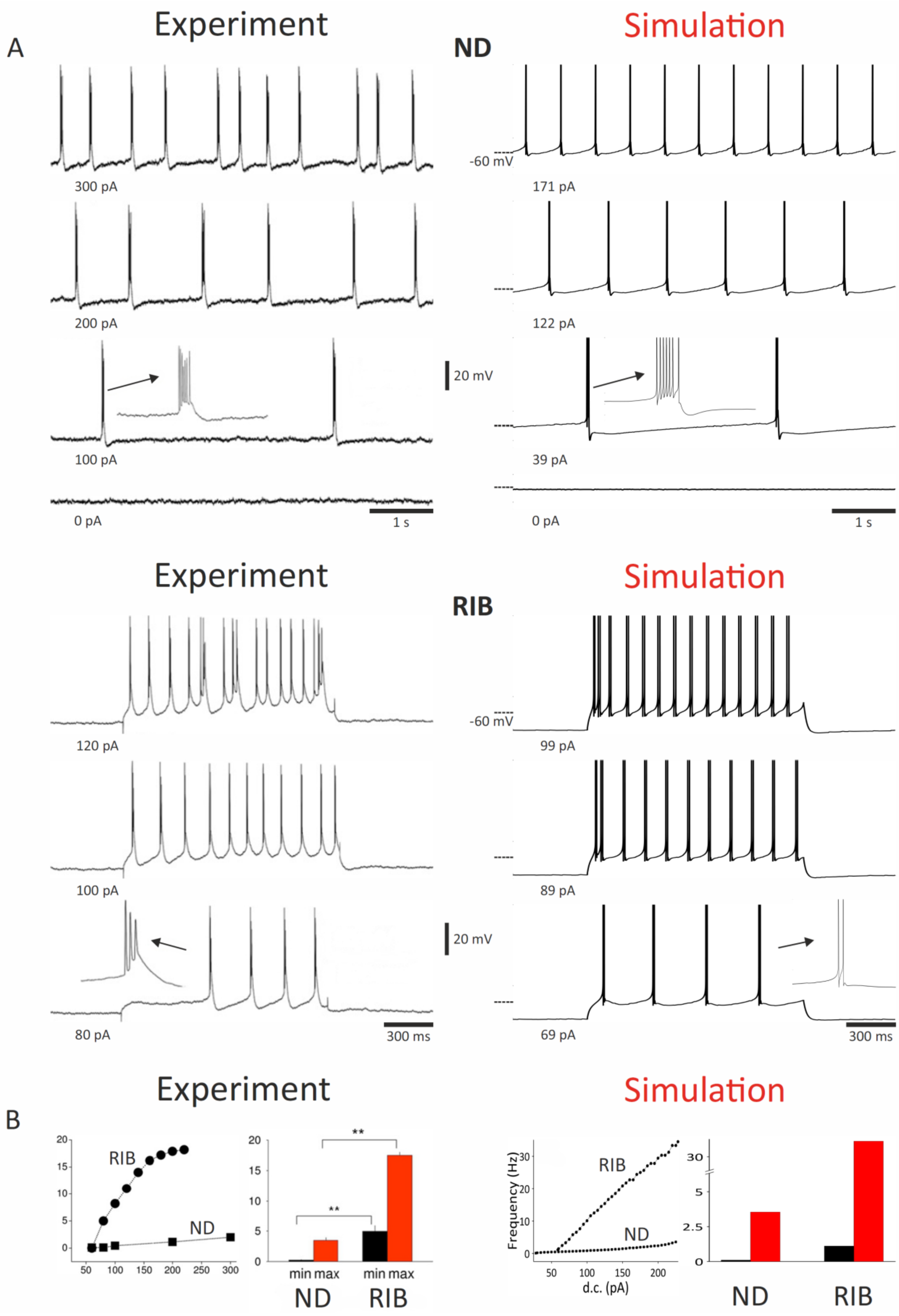

A
Experiment
Simulation
ND
-60 mV
300 pA
171 pA
200 pA
122 pA
20 mV
100 pA
39 pA
0 pA
1 s
0 pA
1 s
Experiment
Simulation
RIB
-60 mV
120 pA
99 pA
100 pA
89 pA
20 mV
80 pA
300 ms
69 pA
300 ms
B
Experiment
Simulation
RIB
ND
**
**
min max
min max
ND
RIB
Frequency (Hz)
RIB
ND
d.c. (pA)
ND
RIB

**Figure S4. Experimental and simulated firing patterns of ND and RIB neurons.**

A, For each neuron type, intracellularly recorded firing pattern from the mouse cortex (Experiment) and simulated activity (Simulation). Value of injected current is indicated below each trace. B, Plots of frequency versus injected current and histograms of minimal and maximal bursting frequencies for ND and RIB neurons as observed in the experiments and the simulations. Dashed line on the left of the traces indicates -60 mV. Experimental data are reproduced with permission from Lorincz, Gunner, Bao, Connelly, Isaac, Hughes and Crunelli (2015)[2].

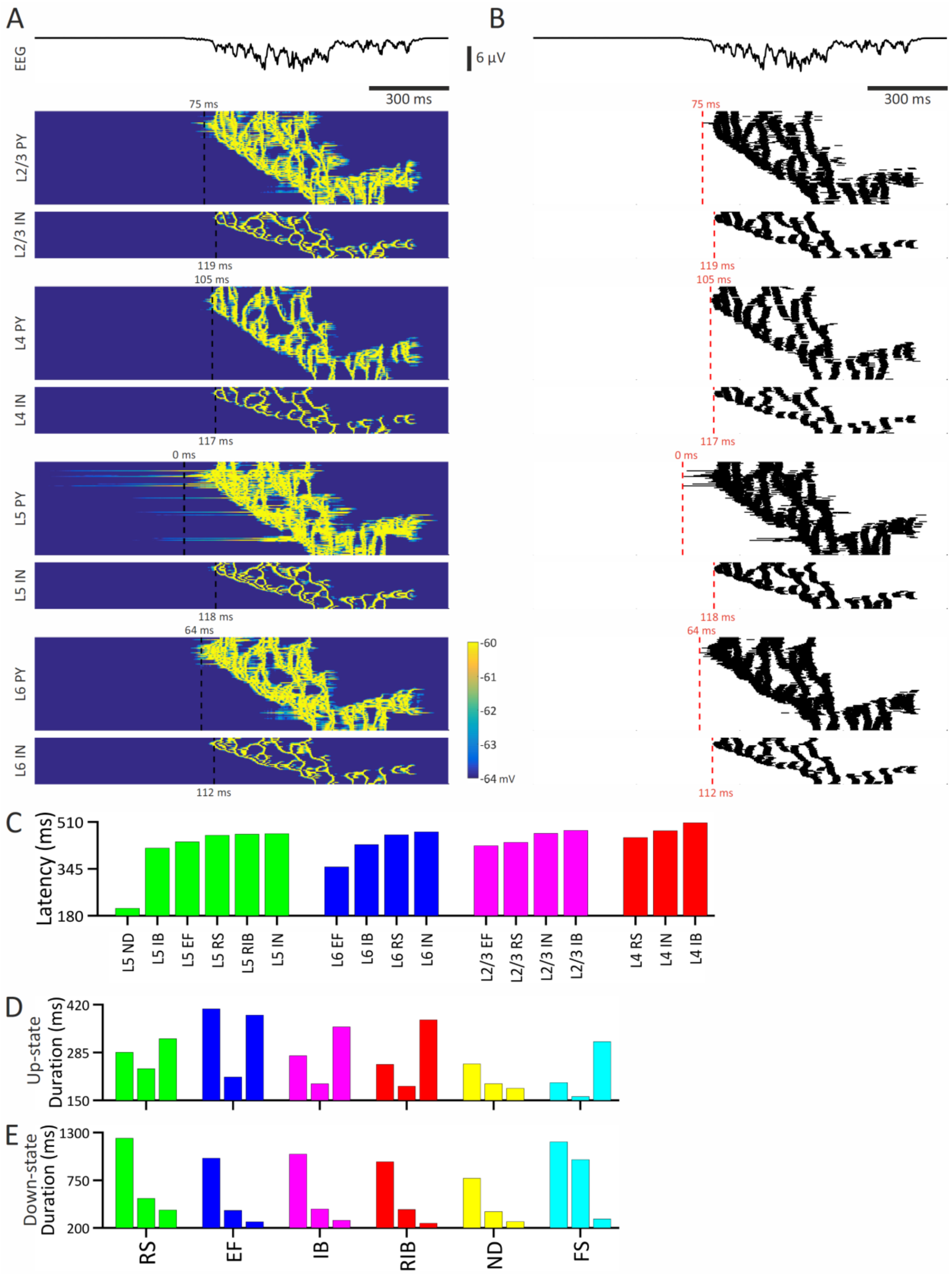
A
B
EEG
6 µV
300 ms
75 ms
L2/3 PY
L2/3 IN
119 ms
105 ms
L4 PY
L4 IN
117 ms
0 ms
L5 PY
L5 IN
118 ms
64 ms
L6 PY
L6 IN
112 ms
-60
-61
-62
-63
-64 mV
C
Latency (ms)
510
345
180
L5 ND
L5 IB
L5 EF
L5 RS
L5 RIB
L5 IN
L6 EF
L6 IB
L6 RS
L6 IN
L2/3 EF
L2/3 RS
L2/3 IN
L2/3 IB
L4 RS
L4 IN
L4 IB
D
Up-state Duration (ms)
420
285
150
E
Down-state Duration (ms)
1300
750
200
RS
EF
IB
RIB
ND
FS

**Figure S5. Onset timing of firing in simulated Up-states in the isolated cortical network.** A, EEG (top trace) and colour-coded membrane potential graphs of the indicated cortical neuronal populations during a cycle of the slow (< 1Hz) oscillation. B, EEG (top trace) and AP rastergrams corresponding to the colour-coded graphs of the neuronal populations indicated in A. Red dashed vertical line represents the first AP of the Up-state in each population. The latency (indicated in red below each rastergram) is measured relatively to the first AP of the cycle in a layer 5 pyramidal neurons (time zero). C, Mean onset latency histograms of the first AP for all cortical neurons grouped by their type measured with respect to the first AP of an Up-state. D and E, Up- and Down-state durations, respectively, averaged over a 300 sec-long simulation.

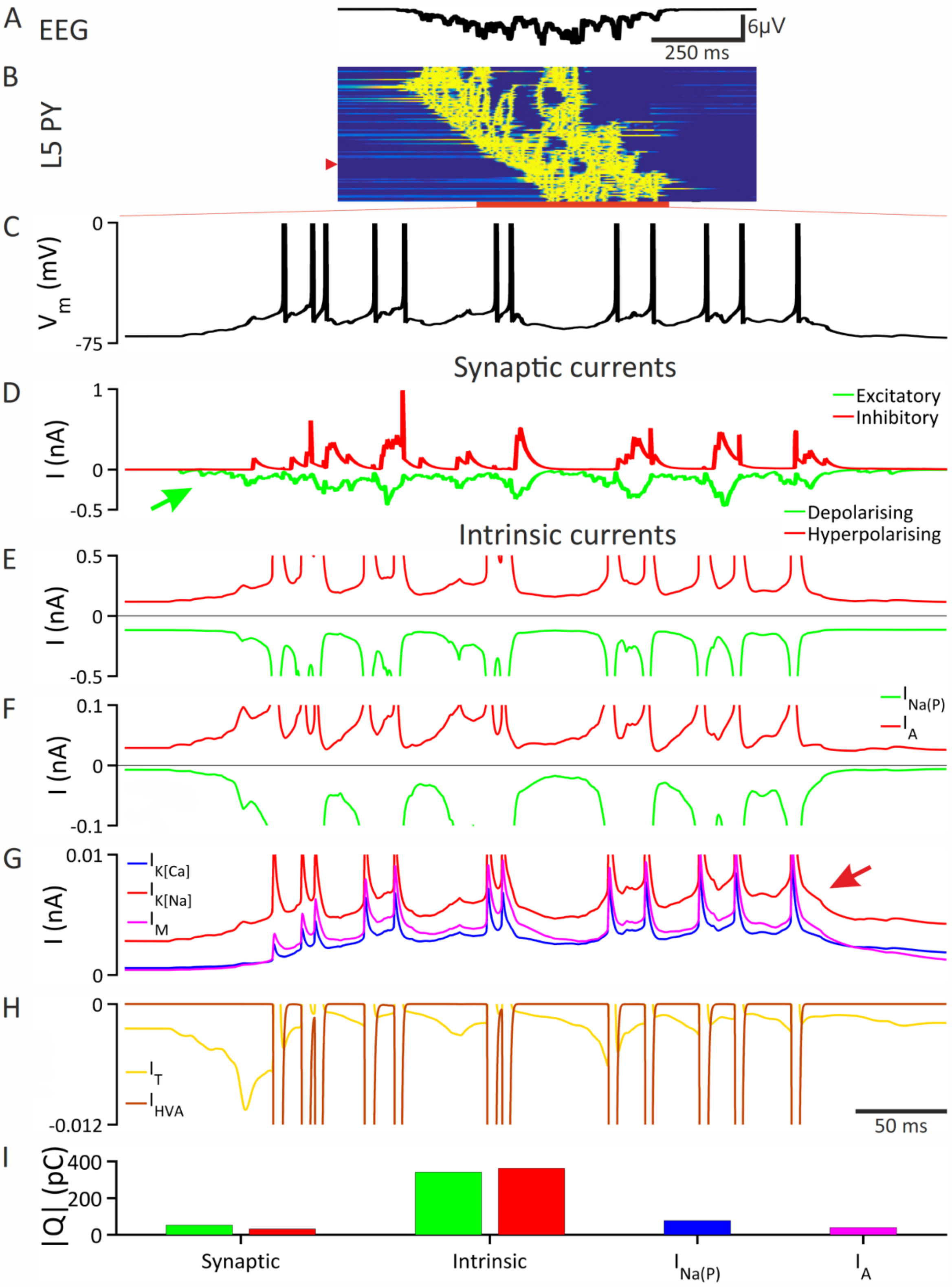

A
EEG
6μV
250 ms
B
L5 PY
C
0
$V_m$ (mV)
-75
Synaptic currents
D
1
I (nA)
0
-0.5
Excitatory
Inhibitory
Depolarising
Hyperpolarising
Intrinsic currents
E
0.5
I (nA)
0
-0.5
F
0.1
I (nA)
0
-0.1
$I_{Na(P)}$
$I_A$
G
0.01
I (nA)
0
$I_{K[Ca]}$
$I_{K[Na]}$
$I_M$
H
0
-0.012
$I_T$
$I_{HVA}$
50 ms
I
|Q| (pC)
400
200
0
Synaptic
Intrinsic
$I_{Na(P)}$
$I_A$

**Figure S6. Membrane currents involved in generating Up-states in the IB neuron model.** A, Simulated EEG showing an UP-state of the slow (< 1Hz) oscillation. B, Colour-coded membrane potential graph of layer 5 pyramidal neurons (L5 PY) during the UP-state shown in A. C, Intracellularly recorded Up-state of the IB neuron marked by the red arrowhead in B. D, Total excitatory and inhibitory synaptic currents during the Up-state shown in C. The green arrow points to EPSPs initiating the Up-state. E, Total current of all intrinsic membrane channels during the Up-state shown in C. E-G, Specific intrinsic membrane currents as indicated in each panel during the Up-state shown in C ($I_{Na(P)}$: persistent $Na^+$ current; $I_A$: A current; $I_{K[Ca]}$: $Ca^{2+}$-activated $K^+$ current; $I_{K[Na]}$, $Na^+$-activated $K^+$ current; $I_M$: M current; $I_T$: T-type $Ca^{2+}$ current; $I_{HVA}$: high-threshold $Ca^{2+}$ current). Red arrow in G points to the accumulation of $K^+$ currents that contributes to the termination of the Up-state. Some currents are truncated for clarity. I, Plots of the indicated total currents during the Up-state shown in C.

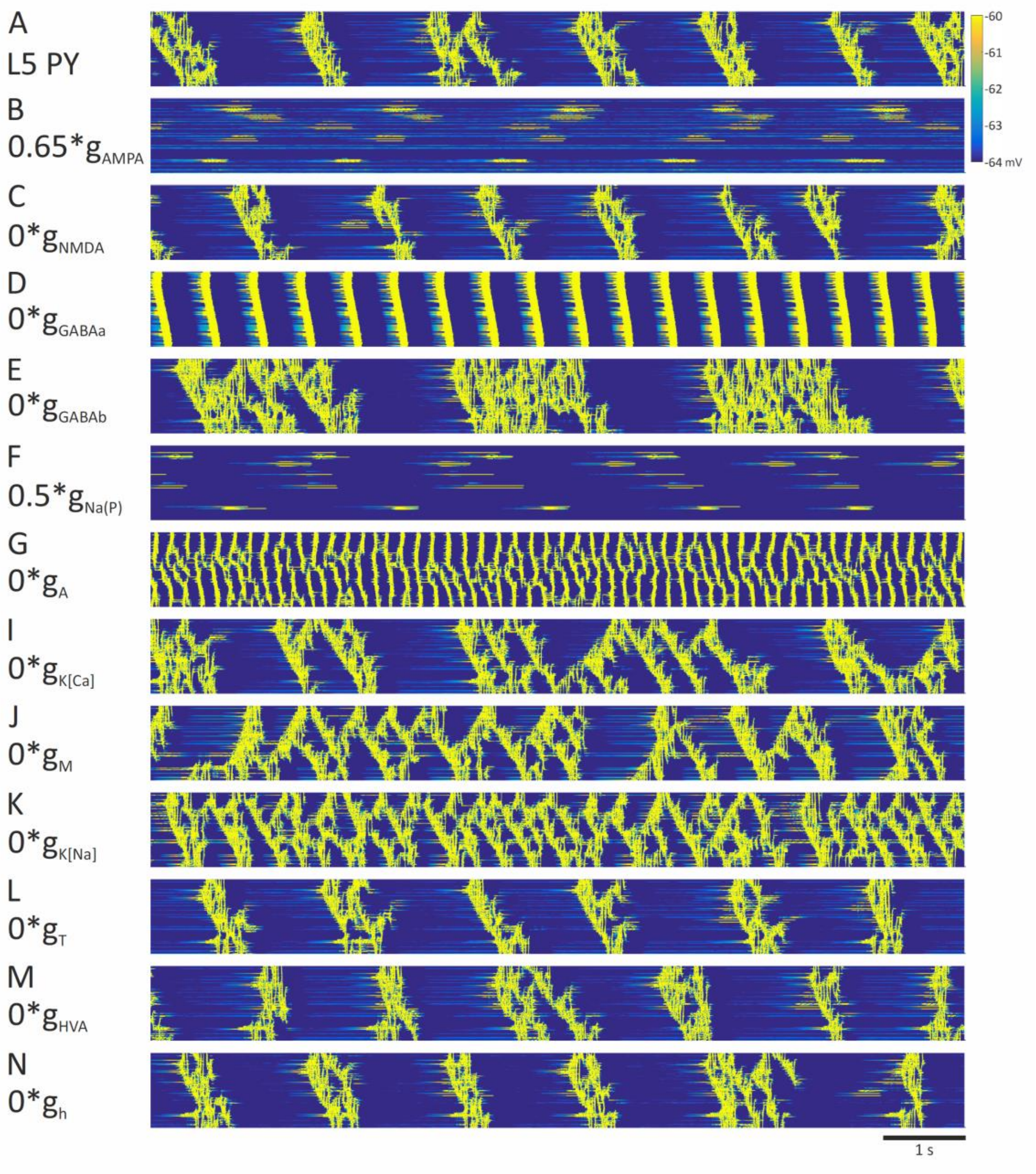


**Figure S7. Role of various synaptic and intrinsic membrane currents in generating Up-states in the isolated neocortical network.**

A, colour-coded membrane potential graphs of the pyramidal neuron population in layer 5 (L5 PY) during simulated slow (<1 Hz) oscillations in the isolated cortical network. B-N, As in A but for simulations obtained following the manipulation of the conductances (g) indicated to the left of each graph ($g_{AMPA}$: AMPA receptor conductance; $g_{NMDA}$: NMDA receptor conductance; $g_{GABAa}$: GABA-A receptor conductance; $g_{GABAb}$: GABA-B receptor conductance; $g_{Na(P)}$: persistent $Na^+$ current conductance; $g_A$: A current conducatnce; $g_{K(Ca)}$: $Ca^{2+}$-activated $K^+$ conductance; $g_M$: M current conductance; $g_{K(Na)}$: $Na^+$-activated $K^+$ conductance; $g_T$: T-type $Ca^{2+}$ conductance; $g_{HVA}$: high-voltage $Ca^{2+}$ conductance; $g_h$: hyperpolarization-activated cyclic nucleotide-gated conductance.

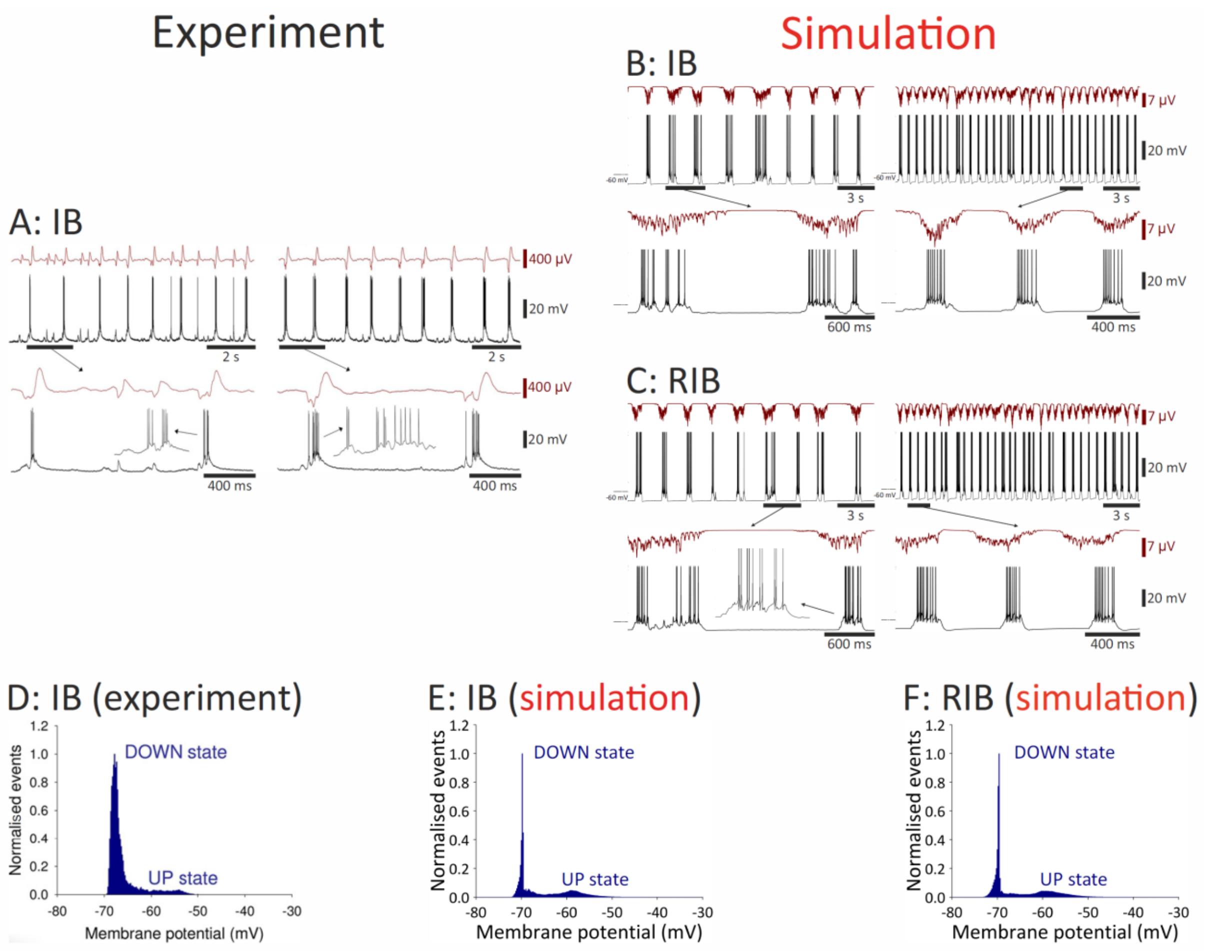


**Figure S8. Experimental and simulated membrane potential dynamics of IB and RIB neurons during slow (<1 Hz) oscillations in the isolated cortical network.**

A,B, Local field potential (top trace) and membrane potential dynamics of an IB neuron recorded *in vitro* (Experiment) and in the isolated cortical network (Simulation) during slow (<1 Hz) oscillations. C, Local field potential (top trace) and membrane potential dynamics of an RIB neuron during simulated slow (<1 Hz) oscillations (Simulation). In A, B and C, the left-hand traces in each pair were recorded during an early appearance of the slow (<1 Hz) oscillation while the right-hand traces show the oscillation at a later stage. D, E, F, Histograms of the normalized membrane potential distribution with the typical peaks of the Up- and Down-state for the experimental and simulated data of an IB neuron and for the simulated results of an RIB neuron. Experimental data are reproduced with permission from Lorincz, Gunner, Bao, Connelly, Isaac, Hughes and Crunelli (2015)[2].

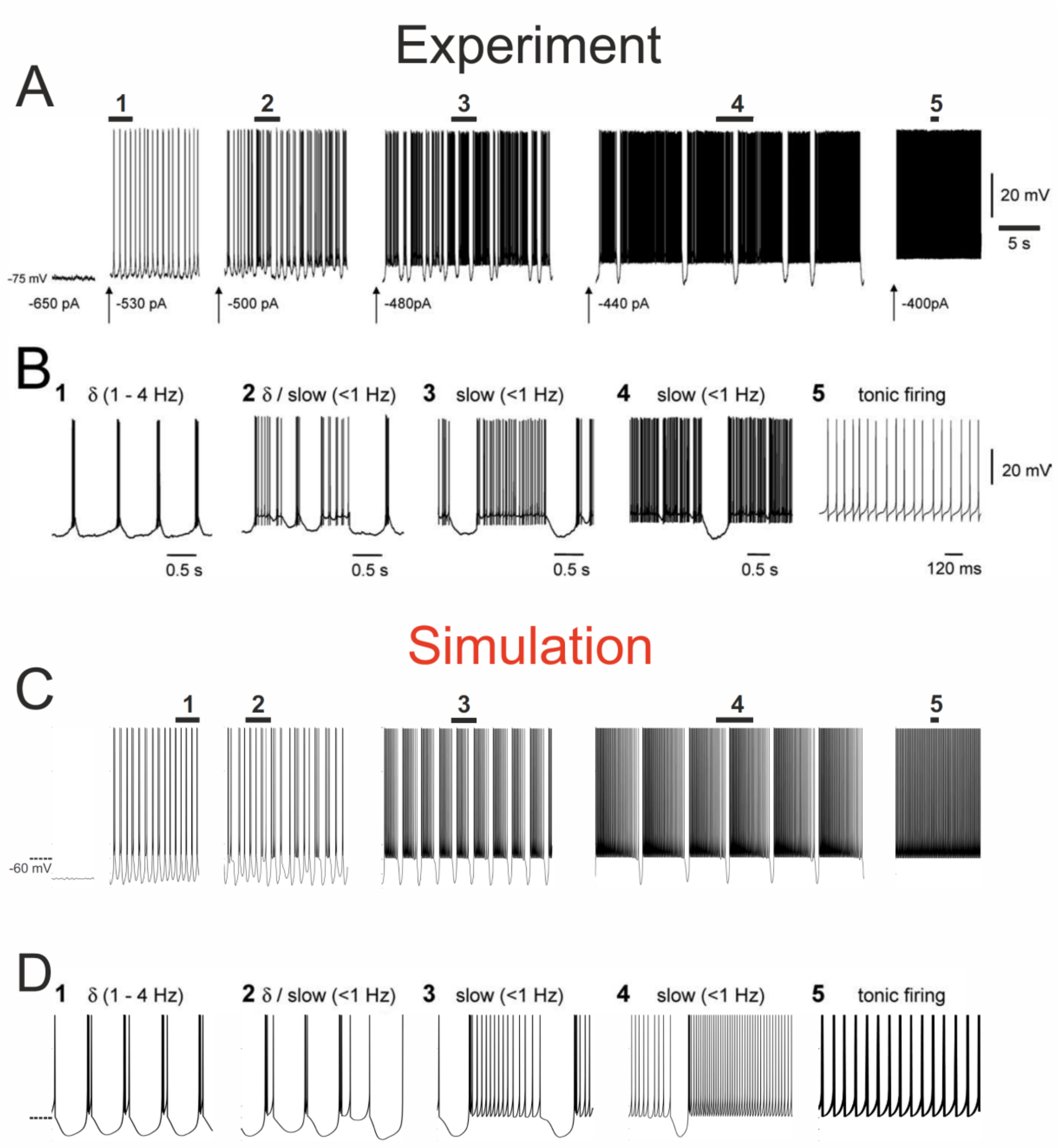


**Figure S9. Intrinsic activity of NRT neurons.**

A, Delta waves, slow (< 1Hz) oscillations and tonic firing of a cat NRT neuron recorded *in vitro*. B, Enlargement of the traces marked by numbers in A. C, Simulated delta waves, slow (< 1Hz) oscillations and tonic firing of an NRT neuron. D, Enlargement of the trace marked by numbers in C. Experimental data are reproduced with permission from Blethyn, Hughes, Tóth, Cope and Crunelli (2006)[3].

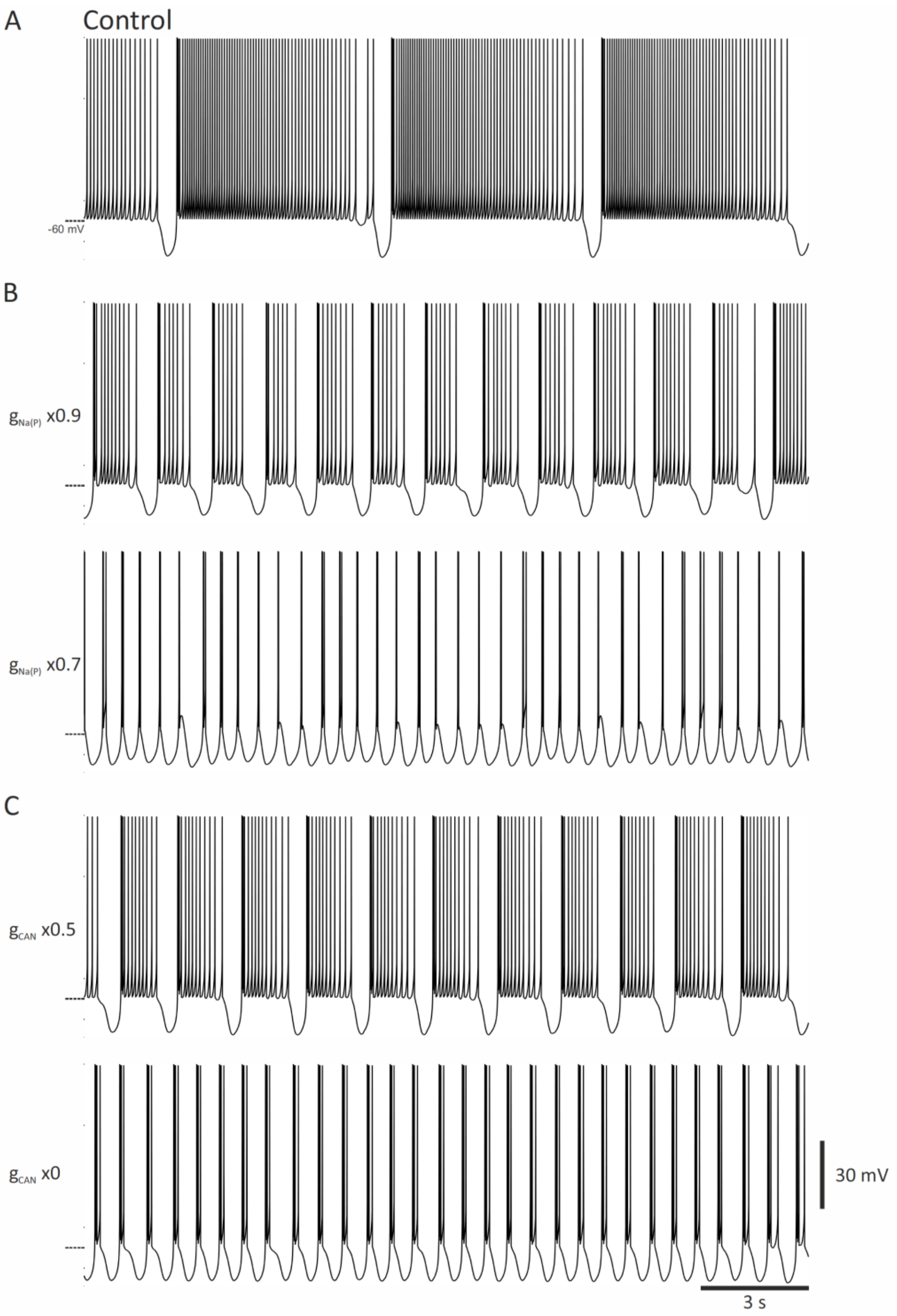


**Figure S10. Role of $I_{Na(P)}$ and $I_{CAN}$ in the slow (< 1Hz) oscillation of $NRT_{FO}$ neurons.**
A, Simulated slow (< 1Hz) oscillations of an $NRT_{FO}$ neuron under control condition. B, Reducing the conductance of the persistent $Na^+$ current ($g_{Na(P)}$) gradually reduces the Up-state duration of the slow (< 1Hz) oscillation, eventually leading to the expression of delta waves (bottom trace). C, Reducing and blocking the conductance of the non-selective cation current ($g_{CAN}$) increases the frequency of the simulated slow (< 1Hz) oscillation and eventually abolishes it, leading to delta waves (bottom trace).

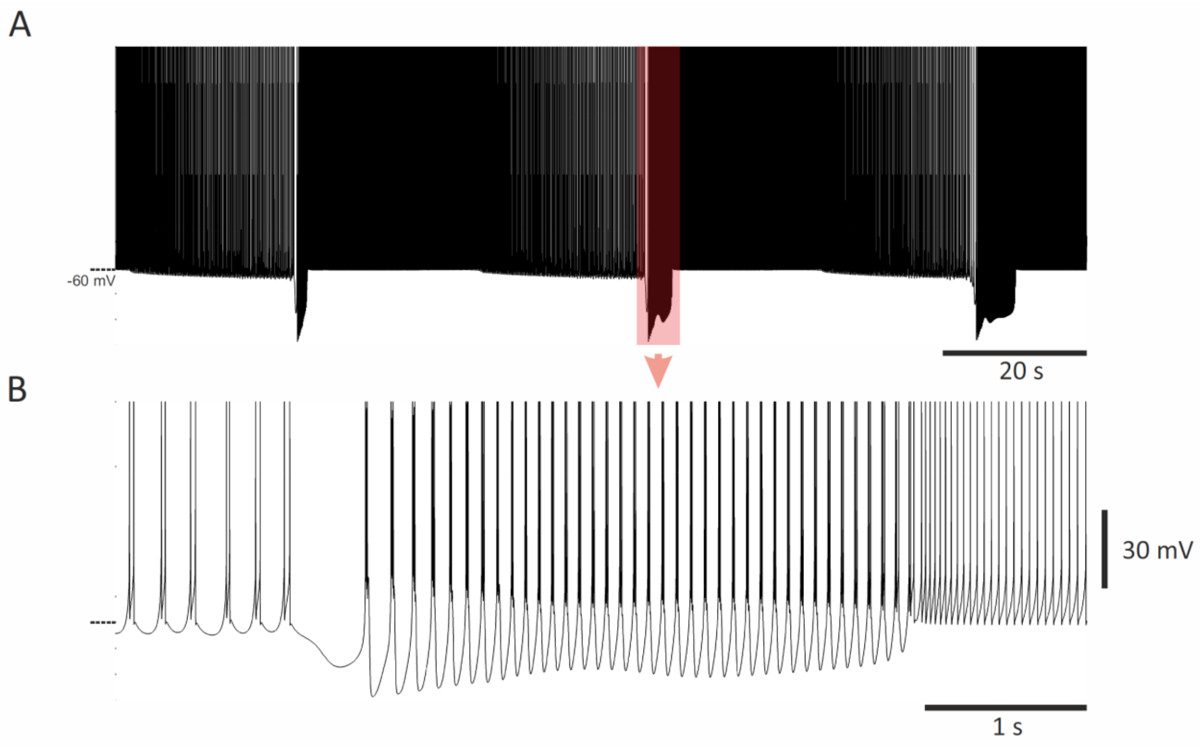


**Figure S11. Spindle waves in isolated NRT$_{FO}$ neurons.**

A, The trace shows 3 fast (~9-12 Hz) spindle waves that were generated spontaneously in NRT$_{FO}$ neurons when they are isolated from TC$_{FO}$ neurons. B, Enlargement of the spindle wave highlight in A.

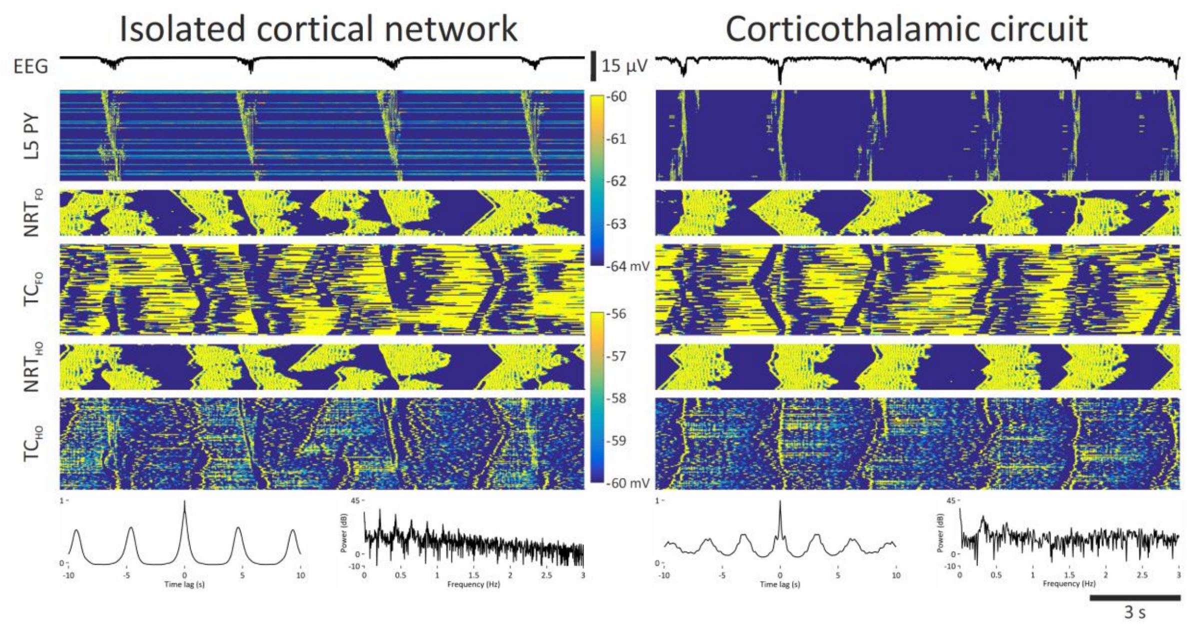


**Figure S12. The thalamic input increases the frequency of the slow (<1 Hz) oscillation.** Left panel: simultaneous EEG (top trace) and colour-coded membrane potential graphs of the indicated neuronal populations during slow (<1 Hz) oscillations simulated in the isolated cortical model. EEG autocorrelograms (left) and power graphs (right) are shown at the bottom. Right panel: as on the left but with an active thalamocortical input. Note the increased regularity and frequency of the slow (< 1Hz) oscillation in B compared to that shown in A. L5 PY: pyramidal neurons in cortical layer 5; $NRT_{FO}$: first order NRT neurons; $TC_{FO}$: first order TC neurons; $NRT_{HO}$: higher order NRT neurons; $TC_{HO}$: higher order TC neurons.

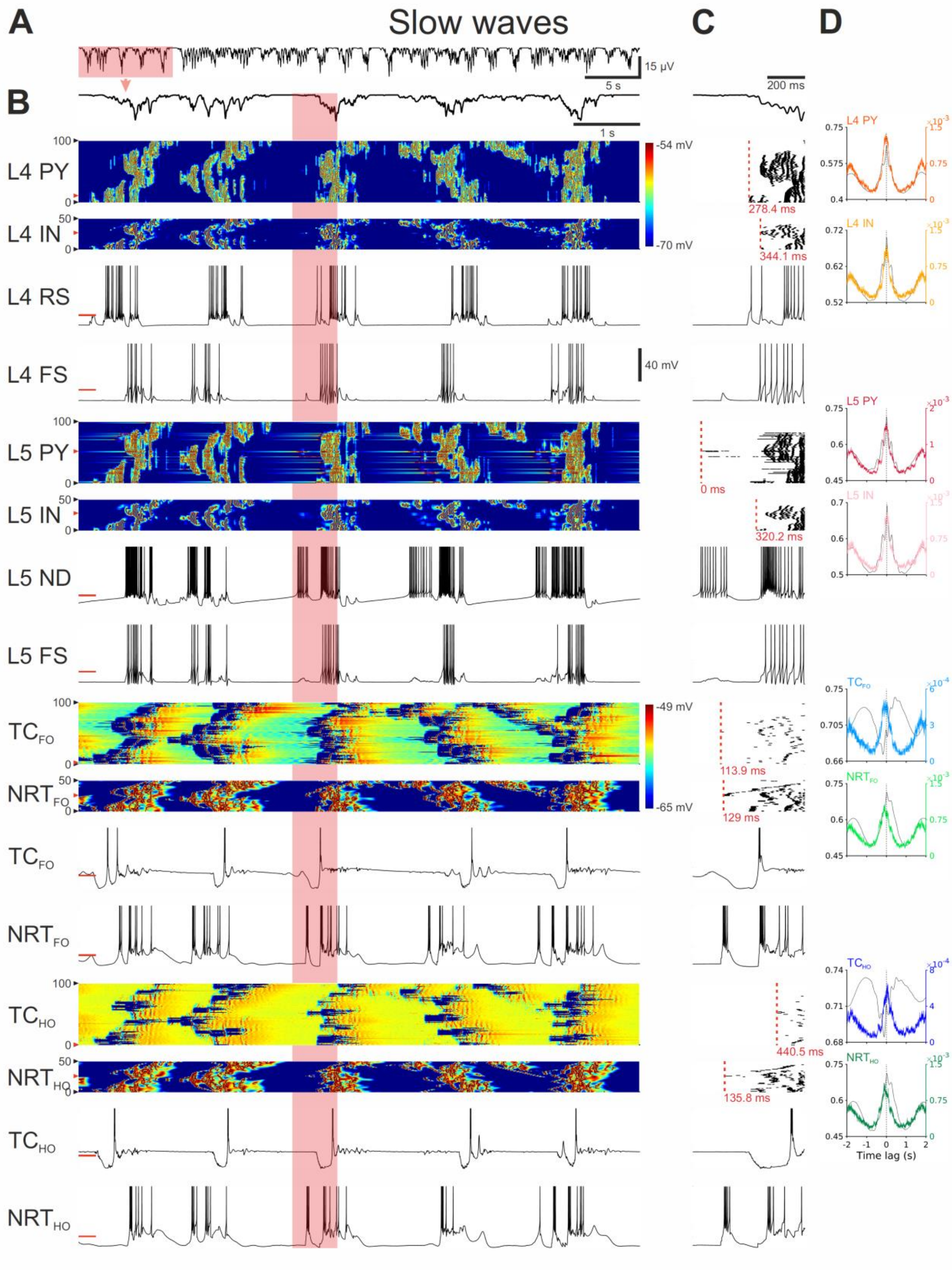

A
Slow waves
C
D
15 µV
5 s
B
1 s
200 ms
L4 PY
L4 IN
L4 RS
L4 FS
L5 PY
L5 IN
L5 ND
L5 FS
TC FO
NRT FO
TC FO
NRT FO
TC HO
NRT HO
TC HO
NRT HO
-54 mV
-70 mV
40 mV
-49 mV
-65 mV
278.4 ms
344.1 ms
0 ms
320.2 ms
113.9 ms
129 ms
440.5 ms
135.8 ms
Time lag (s)

**Figure S13. Slow (<1Hz) oscillations in the full corticothalamic model start in L5 when ND neurons are depolarized.**

A, EEG showing the rhythmic pattern of slow (< 1Hz) oscillations. B, EEG (top trace) and colour-coded membrane potential plots of the indicated cortical and thalamic neuronal populations during the 5 cycles of the slow (< 1Hz) oscillation highlighted in A (note the two separate colour-scales for the cortical and thalamic neurons). Below are the corresponding membrane potential waveforms of the two neurons indicated by the red arrow on the left of the corresponding colour-coded plots. C, EEG (top trace) and AP rastergrams of the firing in each neuronal population for the slow (< 1Hz) oscillation cycle highlighted in B. Red dashed vertical line represents the first AP of the Up-state in each population. The latency (indicated in red below each rastergram) is measured relatively to the first AP of the cycle in the L5 neuron that fires first (time zero). Below the rastergrams are the corresponding membrane potential waveforms of that cycle for the indicated neuron. D, Cross-correlations of EEG and APs for the indicated neuronal populations, calculated over a 485 sec-long simulation. Shaded regions are 95% confidence intervals. Dashed vertical line indicates the zero lag. L4 PY: pyramidal neurons in cortical layer 4; L4 IN: interneurons in cortical layer 4; L4 IB: IB neuron in cortical layer 4; L4 FS: FS neuron in cortical layer 4; L5 PY: pyramidal neurons in cortical layer 5; L5 IN: interneurons in cortical layer 5; L5 RS: RS neuron in cortical layer 5; L5 FS: FS neuron in cortical layer 5; $TC_{FO}$: first order TC neurons; $TC_{HO}$: higher order TC neurons; $NRT_{FO}$: first order NRT neurons; $NRT_{HO}$: higher order NRT neurons.

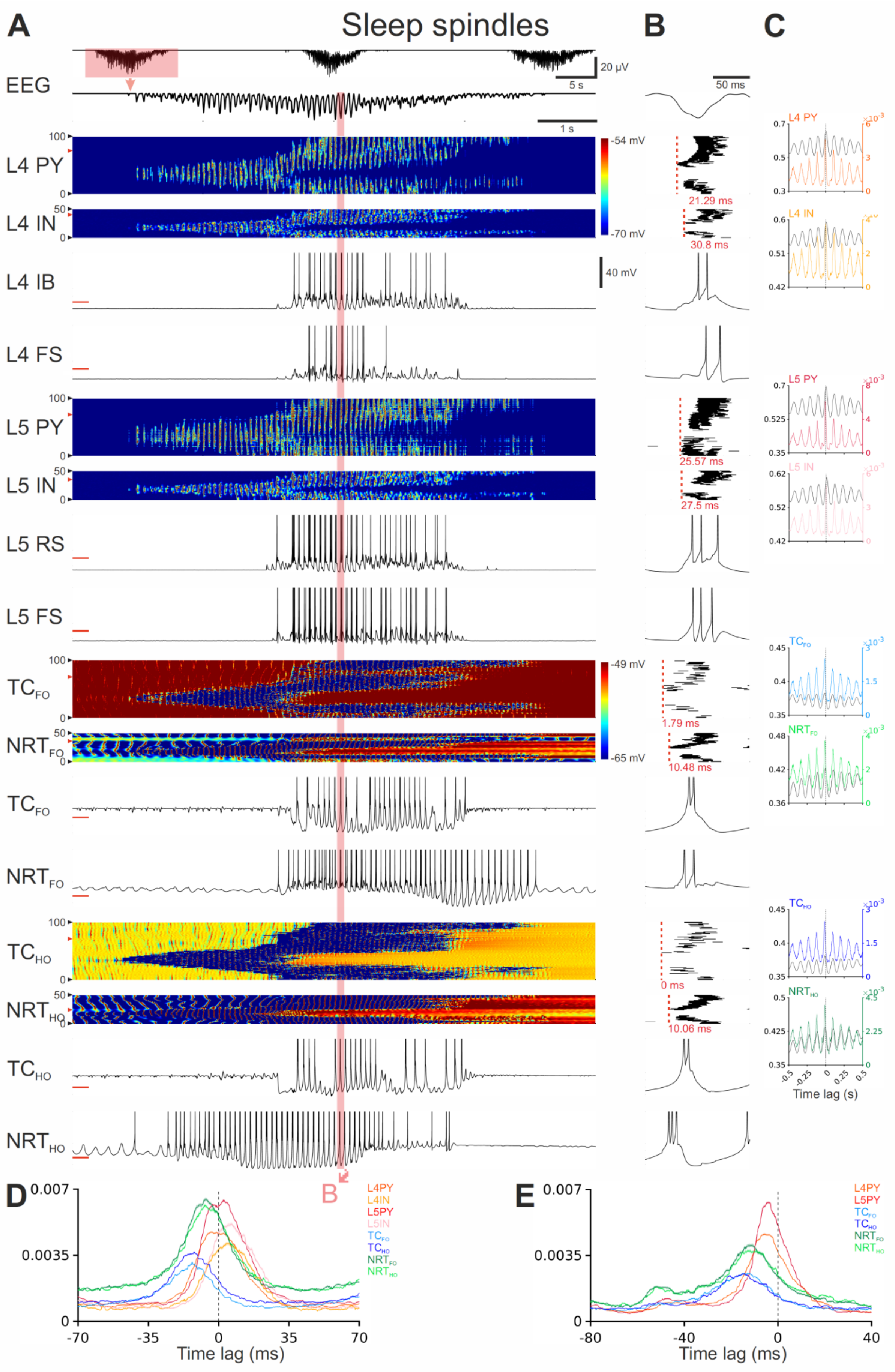

A
Sleep spindles
B
C
EEG
20 µV
5 s
50 ms
1 s
L4 PY
L4 IN
L4 IB
L4 FS
L5 PY
L5 IN
L5 RS
L5 FS
TCFO
NRTFO
TCHO
NRTHO
-54 mV
-70 mV
40 mV
-49 mV
-65 mV
21.29 ms
30.8 ms
25.57 ms
27.5 ms
1.79 ms
10.48 ms
0 ms
10.06 ms
Time lag (s)
D
E
Time lag (ms)
L4PY
L4IN
L5PY
L5IN
TCFO
TCHO
NRTFO
NRTHO

**Figure S14. Sleep spindles in the full corticothalamic model.**

A, Top trace: EEG showing the rhythmic pattern of sleep spindles. The highlighted spindle wave is enlarged below. Lower traces: colour-coded membrane potential plots of the indicated cortical and thalamic neuron populations (note the two separate colour-coded scales for the cortex and the thalamus). Below are the corresponding membrane potential waveforms of the two neurons indicated by the red arrow on the left of the corresponding colour-coded plots. B, EEG (top trace), AP rastergrams of the firing of the first AP in each neuronal population for the sleep spindle cycle highlighted in B. Red dashed vertical line represents the first AP of the Up-state in each population. The latency (indicated below each rastergram) is measured relatively to the first AP of the cycle in a $TC_{HO}$ neuron (time zero). Below the rastergrams are the corresponding membrane potential waveforms of that cycle for the indicated neuron. C, Cross-correlations of EEG and APs for the indicated neuronal populations, calculated over a 485 sec-long simulation. Shaded regions are 95% confidence intervals. Dashed vertical line indicates the zero lag. D, AP distribution with respect to the EEG for all APs of the indicated neuronal populations. Shaded regions are 95% confidence intervals. Dashed vertical line indicates the zero lag. E, Distribution of the first AP in a spindle cycle with respect to the EEG for all APs of the indicated neuronal populations. Shaded regions are 95% confidence intervals. Dashed vertical line indicates the zero lag. L4 PY: pyramidal neurons in cortical layer 4; L4 IN: interneurons in cortical layer 4; L4 IB: IB neuron in cortical layer 4; L4 FS: FS neuron in cortical layer 4; L5 PY: pyramidal neurons in cortical layer 5; L5 IN: interneurons in cortical layer 5; L5 RS: RS neuron in cortical layer 5; L5 FS: FS neuron in cortical layer 5; $TC_{FO}$: first order TC neurons; $TC_{HO}$: higher order TC neurons; $NRT_{FO}$: first order NRT neurons; $NRT_{HO}$: higher order NRT neurons.

### Supplementary Tables

**Table S1. Network connectivity parameters.**

| Grp | Src | Target | Type | P (%) | Amp (mV), weight | Mini (mV), weight | Del (ms) | RT (ms) | $\tau_D$ (ms) |
|---|---|---|---|---|---|---|---|---|---|
| 1.1 | $L2/3_E$ | $L2/3_E$ | AMPA | 13 | 0.8, 1.05 | 0.17, 0.23 | 3 | 4.4 | 20 |
| | | | NMDA | 13 | 0.06*, 1.05 | - | 3 | 13.2 | 135 |
| 1.2 | $L2/3_E$ | $L2/3_I$ | AMPA | 13 | 0.8, 1.05 | 0.17, 0.17 | 1.5 | 4.4 | 22 |
| | | | NMDA | 13 | 0.06*, 1.05 | - | 1.5 | 13.2 | 90 |
| 1.3 | $L2/3_I$ | $L2/3_E$ | $GABA_A$ | 13 | 1-1.5, 2.25 | 0.083, 0.083 | 1.5 | 3.5 | 22 |
| | | | $GABA_B$ | 13 | 4.25**, 2.25 | - | 1.5 | 42.5 | 90 |
| 2.1 | $L2/3_E$ | $L4_E$ | AMPA | 3 | 0.8, 1.05 | 0.17, 0.23 | 2.5 | 4.4 | 20 |
| | | | NMDA | 3 | 0.06*, 1.05 | - | 2.5 | 13.2 | 135 |
| 2.2 | $L2/3_E$ | $L4_I$ | $GABA_A$ | 3 | 1-1.5, 2.25 | 0.17, 0.17 | 2.5 | 3.5 | 22 |
| | | | $GABA_B$ | 3 | 4.25**, 2.25 | - | 2.5 | 42.5 | 90 |
| 3.1 | $L2/3_E$ | $L5_E$ | AMPA | 20 | 0.8, 1.05 | 0.17, 0.23 | 2.75 | 4.4 | 20 |
| | | | NMDA | 20 | 0.06*, 1.05 | - | 2.75 | 13.2 | 135 |
| 3.2 | $L2/3_E$ | $L5_I$ | $GABA_A$ | 20 | 1-1.5, 2.25 | 0.17, 0.17 | 2.75 | 3.5 | 22 |
| | | | $GABA_B$ | 20 | 4.25**, 2.25 | - | 2.75 | 42.5 | 90 |
| 4.1 | $L4_E$ | $L4_E$ | AMPA | 15 | 0.8 1.05 | 0.17, 0.23 | 3 | 4.4 | 20 |
| | | | NMDA | 15 | 0.06* 1.05 | - | 3 | 13.2 | 135 |
| 4.2 | $L4_E$ | $L4_I$ | AMPA | 15 | 0.8 1.05 | 0.17, 0.17 | 1.5 | 4.4 | 20 |
| | | | NMDA | 15 | 0.06* 1.05 | - | 1.5 | 13.2 | 135 |
| 4.3 | $L4_I$ | $L4_E$ | $GABA_A$ | 15 | 1-1.5, 2.25 | 0.083, 0.083 | 1.5 | 3.5 | 22 |
| | | | $GABA_B$ | 15 | 4.25**, 2.25 | - | 1.5 | 42.5 | 90 |
| 5.1 | $L4_E$ | $L2/3_E$ | AMPA | 25 | 0.8, 1.05 | 0.17, 0.23 | 2.5 | 4.4 | 20 |
| | | | NMDA | 25 | 0.06*, 1.05 | - | 2.5 | 13.2 | 135 |
| 5.2 | $L4_E$ | $L2/3_I$ | $GABA_A$ | 25 | 1-1.5, 2.25 | 0.17, 0.17 | 2.5 | 3.5 | 22 |
| | | | $GABA_B$ | 25 | 4.25**, 2.25 | - | 2.5 | 42.5 | 90 |
| 6.1 | $L4_E$ | $L5_E$ | AMPA | 9 | 0.8, 1.05 | 0.17, 0.23 | 2.5 | 4.4 | 20 |
| | | | NMDA | 9 | 0.06*, 1.05 | - | 2.5 | 13.2 | 135 |
| 6.2 | $L4_E$ | $L5_I$ | $GABA_A$ | 9 | 1-1.5, 2.25 | 0.17, 0.17 | 2.5 | 3.5 | 22 |
| | | | $GABA_B$ | 9 | 4.25**, 2.25 | - | 2.5 | 42.5 | 90 |
| 7.1 | $L4_E$ | $L6_E$ | AMPA | 9 | 0.8, 1.05 | 0.17, 0.23 | 2.75 | 4.4 | 20 |
| | | | NMDA | 9 | 0.06*, 1.05 | - | 2.75 | 13.2 | 135 |
| 7.2 | $L4_E$ | $L6_I$ | $GABA_A$ | 9 | 1-1.5, 2.25 | 0.17, 0.17 | 2.75 | 3.5 | 22 |
| | | | $GABA_B$ | 9 | 4.25**, 2.25 | - | 2.75 | 42.5 | 90 |
| 8.1 | $L5_E$ | $L5_E$ | AMPA | 10 | 0.8, 1.05 | 0.17, 0.23 | 3 | 4.4 | 20 |
| | | | NMDA | 10 | 0.06*, 1.05 | - | 3 | 13.2 | 135 |
| 8.2 | $L5_E$ | $L5_I$ | AMPA | 10 | 0.8, 1.05 | 0.17, 0.17 | 1.5 | 4.4 | 20 |
| | | | NMDA | 10 | 0.06*, 1.05 | - | 1.5 | 13.2 | 135 |
| 8.3 | $L5_I$ | $L5_E$ | $GABA_A$ | 10 | 1-1.5, 2.25 | 0.083, 0.083 | 1.5 | 3.5 | 22 |
| | | | $GABA_B$ | 10 | 4.25**, 2.25 | - | 1.5 | 42.5 | 90 |
| 9.1 | $L5_E$ | $L2/3_E$ | AMPA | 9 | 0.8, 1.05 | 0.17, 0.23 | 2.75 | 4.4 | 20 |
| | | | NMDA | 9 | 0.06*, 1.05 | - | 2.75 | 13.2 | 135 |
| 9.2 | $L5_E$ | $L2/3_I$ | $GABA_A$ | 9 | 1-1.5, 2.25 | 0.17, 0.17 | 2.75 | 3.5 | 22 |
| | | | $GABA_B$ | 9 | 4.25**, 2.25 | - | 2.75 | 42.5 | 90 |

| | | | | | | | | | |
|---|---|---|---|---|---|---|---|---|---|
| 10.1 | $L5_E$ | $L4_E$ | AMPA | 3 | 0.8, 1.05 | 0.17, 0.23 | 2.5 | 4.4 | 20 |
| | | | NMDA | 3 | 0.06*, 1.05 | - | 2.5 | 13.2 | 135 |
| 10.2 | $L5_E$ | $L4_I$ | $GABA_A$ | 3 | 1-1.5, 2.25 | 0.17, 0.17 | 2.5 | 3.5 | 22 |
| | | | $GABA_B$ | 3 | 4.25**, 2.25 | - | 2.5 | 42.5 | 90 |
| 11.1 | $L5_E$ | $L6_E$ | AMPA | 15 | 0.8, 1.05 | 0.17, 0.23 | 2.5 | 4.4 | 20 |
| | | | NMDA | 15 | 0.06*, 1.05 | - | 2.5 | 13.2 | 135 |
| 11.2 | $L5_E$ | $L6_I$ | $GABA_A$ | 15 | 1-1.5, 2.25 | 0.17, 0.17 | 2.5 | 3.5 | 22 |
| | | | $GABA_B$ | 15 | 4.25**, 2.25 | - | 2.5 | 42.5 | 90 |
| 12.1 | $L6_E$ | $L6_E$ | AMPA | 10 | 0.8, 1.05 | 0.17, 0.23 | 3 | 4.4 | 20 |
| | | | NMDA | 10 | 0.06*, 1.05 | - | 3 | 13.2 | 135 |
| 12.2 | $L6_E$ | $L6_I$ | AMPA | 10 | 0.8, 1.05 | 0.17, 0.17 | 1.5 | 4.4 | 20 |
| | | | NMDA | 10 | 0.06*, 1.05 | - | 1.5 | 13.2 | 135 |
| 12.3 | $L6_I$ | $L6_E$ | $GABA_A$ | 10 | 1-1.5, 2.25 | 0.083, 0.083 | 1.5 | 3.5 | 22 |
| | | | $GABA_B$ | 10 | 4.25**, 2.25 | - | 1.5 | 42.5 | 90 |
| 13.1 | $L6_E$ | $L4_E$ | AMPA | 15 | 0.8, 1.05 | 0.17, 0.23 | 3.5 | 4.4 | 20 |
| | | | NMDA | 15 | 0.06*, 1.05 | - | 3.5 | 13.2 | 135 |
| 13.2 | $L6_E$ | $L4_I$ | $GABA_A$ | 15 | 1-1.5, 2.25 | 0.17, 0.17 | 3.5 | 3.5 | 22 |
| | | | $GABA_B$ | 15 | 4.25**, 2.25 | - | 3.5 | 42.5 | 90 |
| 14.1 | $L6_E$ | $L5_E$ | AMPA | 3 | 0.8, 1.05 | 0.17, 0.23 | 3.25 | 4.4 | 20 |
| | | | NMDA | 3 | 0.06*, 1.05 | - | 3.25 | 13.2 | 135 |
| 14.2 | $L6_E$ | $L5_I$ | $GABA_A$ | 3 | 1-1.5, 2.25 | 0.17, 0.17 | 3.25 | 3.5 | 22 |
| | | | $GABA_B$ | 3 | 4.25**, 2.25 | - | 3.25 | 42.5 | 90 |
| 15.1 | $NRT_{FO}$ | $NRT_{FO}$ | $GABA_A$ | 10 | 0.10, 0.028 | 0.25, 0.1 | 1 | 12 | 38 |
| 15.2 | $NRT_{HO}$ | $NRT_{HO}$ | $GABA_A$ | 10 | 0.10, 0.028 | 0.25, 0.1 | 1 | 12 | 38 |
| 16.1 | $NRT_{FO}$ | $TC_{FO}$ | $GABA_A$ | 7.5 | 0.4, 0.6 | 0.5, 0.5 | 2.5 | 2.4 | 30 |
| | | | $GABA_B$ | 7.5 | 0.8**, 0.6 | - | 2.5 | 90 | 70 |
| 16.2 | $NRT_{HO}$ | $TC_{HO}$ | $GABA_A$ | 7.5 | 0.4, 0.6 | 0.5, 0.5 | 2.5 | 2.4 | 30 |
| | | | $GABA_B$ | 7.5 | 0.8**, 0.6 | - | 2.5 | 90 | 70 |
| 17.1 | $NRT_{FO}$ | $TC_{HO}$ | $GABA_A$ | 2.5 | 0.4, 0.6 | 0.5, 0.5 | 2.5 | 2.4 | 30 |
| | | | $GABA_B$ | 2.5 | 0.8**, 0.6 | - | 2.5 | 90 | 70 |
| 17.2 | $NRT_{HO}$ | $TC_{FO}$ | $GABA_A$ | 2.5 | 0.4, 0.6 | 0.5, 0.5 | 2.5 | 2.4 | 30 |
| | | | $GABA_B$ | 2.5 | 0.8**, 0.6 | - | 2.5 | 90 | 70 |
| 18.1 | $TC_{FO}$ | $NRT_{FO}$ | AMPA | 3.75 | 4, 0.44 | 0.5, 0.052 | 1 | 0.6 | 16 |
| | | | NMDA | 3.75 | 0.1*, 0.44 | - | 1 | 13.5 | 75 |
| 18.2 | $TC_{HO}$ | $NRT_{HO}$ | AMPA | 3.75 | 4, 0.44 | 0.5, 0.052 | 1 | 0.6 | 16 |
| | | | NMDA | 3.75 | 0.1*, 0.44 | - | 1 | 13.5 | 75 |
| 19.1 | $TC_{FO}$ | $NRT_{HO}$ | AMPA | 1.25 | 4, 0.44 | 0.5, 0.052 | 1 | 0.6 | 16 |
| | | | NMDA | 1.25 | 0.1*, 0.44 | - | 1 | 13.5 | 75 |
| 19.2 | $TC_{HO}$ | $NRT_{FO}$ | AMPA | 1.25 | 4, 0.44 | 0.5, 0.052 | 1 | 0.6 | 16 |
| | | | NMDA | 1.25 | 0.1*, 0.44 | - | 1 | 13.5 | 75 |
| 20.1 | $TC_{FO}$ | $L4_E$ | AMPA | 16 | 5.5, 8.4 | 0.17, 0.17 | 4 | 4.3 | 20.4 |
| | | | NMDA | 16 | 0.4*, 8.4 | - | 4 | 13.6 | 129 |
| 20.2 | $TC_{FO}$ | $L4_I$ | AMPA | 4 | 7, 8.4 | 0.17, 0.17 | 4 | 3.9 | 18.8 |
| | | | NMDA | 4 | 0.5*, 8.4 | - | 4 | 12.6 | 137 |
| 21.1 | $TC_{FO}$ | $L5_E$ | AMPA | 16 | 0.5, 0.7 | 0.17, 0.17 | 4 | 5.2 | 19.4 |
| | | | NMDA | 16 | 0.03*, 0.7 | - | 4 | 13 | 138 |
| 21.2 | $TC_{FO}$ | $L5_I$ | AMPA | 4 | 0.65, 0.7 | 0.17, 0.17 | 4 | 4.2 | 20.1 |

| | | | | | | | | | |
|---|---|---|---|---|---|---|---|---|---|
| | | | NMDA | 4 | 0.04*, 0.7 | - | 4 | 13 | 137 |
| 22.1 | $TC_{HO}$ | $L2/3_E$ | AMPA | 32 | 2.4, 3.5 | 0.17, 0.17 | 6 | 4.3 | 20.7 |
| | | | NMDA | 32 | 0.2*, 3.5 | - | 6 | 13.2 | 142 |
| 22.2 | $TC_{HO}$ | $L2/3_I$ | AMPA | 8 | 3.1, 3.5 | 0.17, 0.17 | 6 | 4.3 | 19.3 |
| | | | NMDA | 8 | 0.2*, 3.5 | - | 6 | 12.9 | 139 |
| 23.1 | $TC_{HO}$ | $L4_E$ | AMPA | 32 | 0.5, 0.7 | 0.17, 0.17 | 6 | 5.1 | 20.2 |
| | | | NMDA | 32 | 0.03*, 0.7 | - | 6 | 13.1 | 132 |
| 23.2 | $TC_{HO}$ | $L4_I$ | AMPA | 8 | 0.65, 0.7 | 0.17, 0.17 | 6 | 4.2 | 20.1 |
| | | | NMDA | 8 | 0.04*, 0.7 | - | 6 | 13.7 | 122 |
| 24.1 | $TC_{HO}$ | $L5_E$ | AMPA | 32 | 3.3, 4.9 | 0.17, 0.17 | 6 | 4.5 | 20.8 |
| | | | NMDA | 32 | 0.25*, 4.9 | - | 6 | 14.7 | 131 |
| 24.2 | $TC_{HO}$ | $L5_I$ | AMPA | 8 | 4.3, 4.9 | 0.17, 0.17 | 6 | 4 | 19.1 |
| | | | NMDA | 8 | 0.3*, 4.9 | - | 6 | 12.3 | 139 |
| 25.1 | $TC_{HO}$ | $L6_E$ | AMPA | 32 | 2.4, 3.5 | 0.17, 0.17 | 6 | 4.5 | 20.2 |
| | | | NMDA | 32 | 0.2*, 3.5 | - | 6 | 13.5 | 131 |
| 25.2 | $TC_{HO}$ | $L6_I$ | AMPA | 8 | 3.1, 3.5 | 0.17, 0.17 | 6 | 4.3 | 19.3 |
| | | | NMDA | 8 | 0.2*, 3.5 | - | 6 | 12.9 | 139 |
| 26 | $L5_E$ | $TC_{HO}$ | AMPA | 10 | 0.025†, 0.05 | 0.5, 0.5 | 4 | 1 | 24.1 |
| | | | NMDA | 10 | 0.0035‡, 0.05 | - | 4 | 20.2 | 83.3 |
| 27 | $L6_E$ | $NRT_{FO}$ | AMPA | 10 | 1.3, 0.132 | 0.5, 0.52 | 8 | 0.8 | 16.4 |
| | | | NMDA | 10 | 0.3*, 0.132 | - | 8 | 15 | 80 |
| 28 | $L6_E$ | $TC_{FO}$ | AMPA | 10 | 0.5†, 0.09 | 0.5, 0.5 | 4 | 0.7 | 22.4 |
| | | | NMDA | 10 | 0.014‡, 0.09 | - | 4 | 22 | 74.4 |
| 29 | $L6_E$ | $NRT_{HO}$ | AMPA | 10 | 1.3, 0.132 | 0.5, 0.52 | 8 | 0.8 | 16.4 |
| | | | NMDA | 10 | 0.3*, 0.132 | - | 8 | 15 | 80 |
| 30 | $L6_E$ | $TC_{HO}$ | AMPA | 10 | 0.5†, 0.09 | 0.5, 0.5 | 4 | 0.7 | 22.4 |
| | | | NMDA | 10 | 0.014‡, 0.09 | - | 4 | 22 | 74.4 |

* Estimated at the resting $V_M$ when extracellular $Mg^{2+}$ concentration is set to 0.1 mM.

** Estimated at the resting $V_M$ in response to a train of 10 presynaptic APs at 100 Hz frequency.

† Estimated at $V_M$ = -80 mV to avoid activation of T-type $Ca^{2+}$ channels.

‡ Estimated at $V_M$ = -80 mV and with extracellular $Mg^{2+}$ concentration set to 0.1 mM.

Abbreviations: E, excitatory cell; I, inhibitory; Grp, projection group; Src, source; P, proportion of connected cells in the target structure; Amp, amplitude; Mini, amplitude of a miniature or spontaneous PSP; Del, delay or latency from an AP in the presynaptic cell (when -10 mV threshold is passed) to the initiation of a PSP; RT, rise time; $\tau_D$, the time it takes for a PSP amplitude to decay to Amp/e, where e is the Euler number.

**Table S2. Derivation of network connectivity parameters.**

| Grp | Type | P | Weight | Delay | Shape |
|---|---|---|---|---|---|
| 1-14 | AMPA, NMDA | [4,5,6,7,8,9,10,11,12,13,14,15,16,17,18,19,20,21,22,23]. | | [18,21,24,25,26,27]. | [28,29,30,31,32]. |
| | $GABA_A$ | [4,5,6,7,8,9,10,11,12,13,14,15,16,17,18,19,20,21,22,23]. | | [18,21,24,25,26,27,33]. | [34,35,36,37,38,39,40]. |
| | $GABA_B$ | [4,5,6,7,8,9,10,11,12,13,14,15,16,17,18,19,20,41,42]. | | [18,21,24,25,26,27,33]. | [40]. |
| 15 | $GABA_A$ | [43,44,45]. | [45,46,47,48,49]. | [46,47,48,49,50,51,52,53,54,55,56,57,58]. | |
| 16-17 | $GABA_A$ | [43,44,45,59]. | [47,50,51,60,61,62,63]. | [64,65]. | [47,50,51]. |
| | $GABA_B$ | [43,44,45,59]. | [47,50,51]. | [64,65]. | [46,61]. |
| 18-19 | AMPA, NMDA | | [61,62,65,66]. | | |
| 20-25 | AMPA, NMDA, $GABA_A$, $GABA_B$ | [59,67,68,69,70,71,72,73,74,75,76,77,78,79,80,81,82,83,84,85,86,87,88,89,90]. | | | |
| 26-30 | AMPA NMDA | [59,63,91,92,93,94,95,96,97,98,99,100,101,102,103,104,105,106]. | | | |

**Table S3. The size of active conductances in TC cells**

| Cell | $\bar{g}_{Na}$ | $\bar{g}_{K(DR)}$ | $\bar{g}_{A1}$ | $\bar{g}_{A2}$ | $\bar{g}_{K1}$ | $\bar{g}_{K2}$ | $\bar{g}_{CAN}$ | $\bar{g}_{h}$ | $\bar{g}_{Na(P)}$ | $\bar{g}_{T}$ | $\bar{P}_{HVA}$ |
|---|---|---|---|---|---|---|---|---|---|---|---|
| $TC_{FO}$ | 70 | 70 | 0.242 | 0.16045 | 0.014 | 0.2 | 0.075 | 8.5 | 0.2015 | 2.1 | 0.135 |
| $TC_{HO}$ | 70 | 70 | 0.242 | 0.16045 | 0.014 | 0.2 | 0.075 | 8.5 | 0.2015 | 4.2 | 0.135 |

The maximum conductances ($\bar{g}$) are expressed in μS/cm$^2$, whereas the maximum membrane permeability to $Ca^{2+}$ ($\bar{P}$) is expressed in μm/s.

**Table S4. The size of active conductances in NRT cells.**

| Cell | $\bar{g}_{Na}$ | $\bar{g}_{K(DR)}$ | $\bar{g}_{AHP1}$ | $\bar{g}_{AHP2}$ | $\bar{g}_{K[Na]}$ | $\bar{g}_{CAN}$ | $\bar{g}_{h}$ | $\bar{g}_{Na(P)}$ | $\bar{g}_{T}$ | $\bar{g}_{HVA}$ |
|---|---|---|---|---|---|---|---|---|---|---|
| NRT | 50 | 50 | 0.003-0.3 | 0.0006-0.06 | 0.0002 | 0.0475 | 0.0043 | 0.01612 | 1.4 | 0.2 |

The maximum conductances ($\bar{g}$) are expressed in μS/cm$^2$.

**Table S5. The size of cortical axosomatic and dendritic active conductances**

| Cell | $\bar{g}_{Na}$ | $\bar{g}_{K(DR)}$ | $\bar{g}_{A}$ | $\bar{g}_{M}$ | $\bar{g}_{fAHP}$ | $\bar{g}_{sAHP}$ | $\bar{g}_{h}$ | $\bar{g}_{Na(P)}$ | $\bar{g}_{K[Na]}$ | $\bar{P}_{T}$ | $\bar{g}_{HVA}$ |
|---|---|---|---|---|---|---|---|---|---|---|---|
| RS | 3000 | - | - | - | - | - | - | 0.077 | 0.07 | - | - |
| | 1.5 | 216 | 1.48 | 0.01 | 0.001 | - | 0.02 | 0.077 | 0.07 | 0.1 | 0.001 |
| EF | 3000 | - | - | - | - | - | - | 0.077 | 0.07 | - | - |
| | 1.5 | 216 | 1.48 | 0.01 | 0.001 | - | 0.02 | 0.077 | 0.07 | 0.1 | 0.001 |
| IB | 3000 | - | - | - | - | - | - | 0.077 | 0.07 | - | - |
| | 1.5 | 216 | 1.48 | 0.01 | 0.001 | - | 0.02 | 0.077 | 0.07 | 1 | 0.01 |
| RIB | 3000 | - | - | - | - | - | - | 0.077 | 0.07 | - | - |
| | 1.5 | 216 | 1.48 | 0.01 | 0.001 | - | 0.02 | 0.077 | 0.07 | 1 | 0.01 |
| SIB | 3000 | - | - | - | - | - | - | 0.077 | 0.07 | - | - |
| | 1.5 | 216 | 1.48 | 0.01 | 0.001 | - | 0.02 | 0.077 | 0.07 | 1 | 0.016 |
| ND | 3000 | - | - | - | - | - | - | 0.077 | 0.07 | - | - |
| | 1.5 | 216 | 1.48 | 0.01 | 0.001 | 0.03 | 0.02 | 0.077 | 0.07 | 1 | 0.016 |
| FS | 3000 | - | - | - | - | - | - | - | 0.07 | - | - |
| | 1.5 | 216 | 1.48 | 0.01 | 0.001 | - | - | - | 0.07 | - | 0.001 |

The maximum conductances ($\bar{g}$) are expressed in mS/cm$^2$, whereas the maximum membrane permeability to $Ca^{2+}$ ($\bar{P}$) is expressed in µm/s. The top and bottom values for each neuron type correspond to axosomatic and dendritic compartments, respectively.

**Table S6. Passive properties of model cells.**

| Cell | $V_R$ (mV) | $R_I$ (MΩ) | τ (ms) | $G_{KL}$ (µS/cm$^2$) | $E_{KL}$ (mV) | $G_{NaL}$ (µS/cm$^2$) | $E_{NaL}$ (mV) | $C_m$ (µf/cm$^2$) | L (µm) | d (µm) |
|---|---|---|---|---|---|---|---|---|---|---|
| $TC_{FO}$ | -65 | 160 | 15.5* | 47 | -90 | 9.1 | 10 | 0.88 | 90 | 60 |
| $TC_{HO}$ | -65 | 130 | 19.2* | 51.2 | -90 | 9.1 | 10 | 0.88 | 90 | 60 |
| NRT | -65 | 160 | 13.7 | 89 | -90 | 23.3 | 10 | 0.88 | 63 | 42 |
| RS | -71.86 | 233 | 15 | 29.3 | -90 | 7.8 | 10 | 0.75 | - | - |
| EF | -65 | >300 | 30 | 15 | -90 | 7.8 | 10 | 0.75 | - | - |
| IB | -71.67 | 227 | 14.8 | 29.3 | -90 | 7.8 | 10 | 0.75 | - | - |
| RIB | -71.67 | 222 | 14.6 | 29.3 | -90 | 7.8 | 10 | 0.75 | - | - |
| SIB | -71.65 | 216 | 14.8 | 29.3 | -90 | 7.8 | 10 | 0.75 | - | - |
| ND | -67.5 | >250 | 25 | 29.3 | -90 | 7.8 | 10 | 0.75 | - | - |
| FS | -73 | 179 | 15.1 | 29.3 | -90 | 7.8 | 10 | 0.75 | - | - |

* Estimated at $V_M$ = -80 mV to avoid activation of T-type $Ca^{2+}$ channels.

The apparent input resistance ($R_I$) was estimated by injecting a hyperpolarising 20 pA current at $V_M$ = -60 mV. Abbreviations: $V_R$, resting membrane potential; $R_I$, apparent input resistance; τ, passive membrane time constant; $E_{KL}$, $K^+$ leak current reversal potential; $E_{NaL}$, $Na^+$ leak current reversal potential; L, length; d, diameter.

**Table S7. Passive membrane parameters of cortical cells.**

| Cell | $g_{SD}$ (nS) | $R_{SD}$ (MΩ) | $A_S$ (μm$^2$) | $A_D$ (μm$^2$) | ρ |
|---|---|---|---|---|---|
| RS | 100.75 | 9.93 | 100.07 | 16011.94 | 160 |
| EF | 100.75 | 9.93 | 100.07 | 16011.94 | 160 |
| IB | 97.69 | 10.24 | 100.07 | 16512.31 | 165 |
| RIB | 94.82 | 10.55 | 100.07 | 17012.68 | 170 |
| SIB | 92.11 | 10.86 | 100.07 | 17513.06 | 175 |
| ND | 100.75 | 9.93 | 100.07 | 16011.94 | 160 |
| FS | 134.33 | 7.44 | 100.07 | 12008.95 | 120 |

Abbreviations: $R_{SD}$ is the resistance between the axosomatic and dendritic compartments; ρ is the $A_D/A_S$ ratio.

**Table S8. Leak K$^+$ conductance of cortical cells in cortical network model.**

| Cell | Depolarised | Transitional | Fast slow (delta) | Slow | Hyperpolarised |
|---|---|---|---|---|---|
| RS | 0.0000043 | 0.0000143 | 0.0000173 | 0.0000173 | 0.0000243 |
| EF | 0.0000043 | 0.0000128 | 0.0000128 | 0.0000128 | 0.0000243 |
| IB | 0.0000093 | 0.0000193 | 0.0000223 | 0.0000223 | 0.0000243 |
| RIB | 0.0000093 | 0.0000193 | 0.0000223 | 0.0000223 | 0.0000243 |
| ND | 0.000002 | 0.000002 | 0.000002 | 0.0000143 | 0.0000243 |
| FS | 0.0000043 | 0.0000143 | 0.0000173 | 0.0000173 | 0.0000243 |

Conductances are expressed in mS/cm$^2$.

**Table S9. Leak K$^+$ conductance of all cells in corticothalamic network model.**

| Cell | Depolarised | Slow (L5/TC init) | Slow (TC init) | Delta | Sleep spindles |
|---|---|---|---|---|---|
| RS | 0.0000043 | 0.0000293 | 0.0000293 | 0.0000293 | 0.0000443 |
| EF | 0.0000129 | 0.0000293 | 0.0000293 | 0.0000293 | 0.0000293 |
| IB | 0.0000093 | 0.0000293 | 0.0000293 | 0.0000293 | 0.0000493 |
| RIB | 0.0000093 | 0.0000293 | 0.0000293 | 0.0000293 | 0.0000493 |
| ND | 0.00001396 | 0.0000093 | 0.0000293 | 0.0000293 | 0.0000293 |
| FS | 0.0000093 | 0.0000293 | 0.0000293 | 0.0000293 | 0.0000293 |
| $TC_{FO}$ | 0.00133 | 0.0015 | 0.0015 | 0.0019 | 0.0008 |
| $TC_{HO}$ | 0.0044 | 0.0022 | 0.0022 | 0.00305 | 0.0021 |
| NRT | 0.0002 | 0.0004 | 0.0004 | 0.0009 | 0.0004 |

Conductances are expressed in mS/cm$^2$ in cortical cells in μS in thalamic cells.

**Table S10. Sets of AMPAR parameters used in the corticothalamic network model.**

| Synapse | $\bar{g}$ (μS) | E (mV) | α ($ms^{-1}$) | β ($ms^{-1}$) | $T_{dur}$ (ms) | $T_{max}$ (mM) |
|---|---|---|---|---|---|---|
| Cortex | 0.001945 | 0 | 0.94 | 0.22 | 0.55 | 0.5 |
| NRT | 0.04 | 0 | 10 | 3 | 0.3 | 0.5 |
| TC | 0.034 | 0 | 10 | 3 | 0.3 | 0.5 |

**Table S11. Sets of $GABA_AR$ parameters used in the corticothalamic network model.**

| Synapse | $\bar{g}$ (μS) | E (mV) | α ($ms^{-1}$) | β ($ms^{-1}$) | $T_{dur}$ (ms) | $T_{max}$ (mM) |
|---|---|---|---|---|---|---|
| Cortex | 0.068 | -80 | 0.1 | 0.2 | 0.8 | 0.5 |
| NRT | 0.8 | -70 | 0.01 | 0.04 | 1.5 | 0.5 |
| TC | 1 | -70 | 0.05 | 2 | 1.4 | 0.5 |

**Table S12. Sets of $GABA_BR$ parameters used in the corticothalamic network model.**

| Synapse | $\bar{g}$ (μS) | $k_1$ ($mM^{-1}$ $ms^{-1}$) | $k_2$ ($ms^{-1}$) | $k_3$ ($ms^{-1}$) | $k_4$ ($ms^{-1}$) | $K_d$ ($mM^4$) | $T_{dur}$ (ms) | $T_{max}$ (mM) |
|---|---|---|---|---|---|---|---|---|
| Cortex | 0.001625 | 0.18 | 0.0025 | 0.19 | 0.06 | 17.83 | 0.8 | 0.5 |
| TC | 0.61 | 0.2 | 0.0028 | 0.28 | 0.45 | 100 | 1.4 | 0.5 |

**Table S13. Sets of NMDAR parameters used in the corticothalamic network model.**

| Synapse | $\bar{g}$ (μS) | $Mg_o$ (mM) |
|---|---|---|
| Cortex | 0.00001275 | 2 |
| Cx-NRT | 0.000003 | 0.5 |
| TC-NRT | 0.000003 | 0.5 |
| Cx-TC | 0.00003 | 0.5 |

### Supplementary Methods

*Model neurons*

Thalamic cells were single-compartment Hodgkin-Huxley models described by an equation:

$$C_m \frac{dV_M}{dt} = -G_L(V_M - E_L) - G_{int}(V_M - E_{int}) - G_{syn}(V_M - E_{syn}) - \frac{g_{gap}(V_M - V_N)}{A_M}, \qquad (\boldsymbol{S1})$$

where $C_m$ is the membrane capacitance per unit area in F/cm$^2$, $V_M$ is the membrane potential in mV, $G_L$, $G_{int}$, and $G_{syn}$ are the leak, intrinsic, and synaptic membrane conductances, respectively, in S/cm$^2$, $E_L$, $E_{int}$, and $E_{syn}$ are the reversal potentials for the corresponding conductances in mV, $g_{gap}$ is the gap junction conductance in S, $A_M$ is the membrane area of the cell in cm$^2$, and $V_N$ is the membrane potential of a neighbouring cell (NRT neurons only) connected by a gap junction (mV).

Cortical cells were Hodgkin-Huxley models with separate axosomatic and dendritic compartments. Equations describing the two corresponding compartments were:

$$C_m \frac{dV_S}{dt} = -G_L(V_S - E_L) - G_{int}(V_S - E_{int}) - G_{syn}(V_S - E_{syn}) - \frac{g_{SD}(V_S - V_D)}{A_S}, \qquad (\boldsymbol{S2})$$

$$C_m \frac{dV_D}{dt} = -G_L(V_D - E_L) - G_{int}(V_D - E_{int}) - G_{syn}(V_D - E_{syn}) - \frac{g_{SD}(V_D - V_S)}{A_D}, \qquad (\boldsymbol{S3})$$

where $V_S$ is the axosomatic membrane potential in mV, $V_D$ is the dendritic membrane potential in mV, $g_{SD}$ is the conductance between the two compartments in S, $A_S$ is the membrane area of the axosomatic compartment in cm$^2$, and $A_D$ is the membrane area of the dendritic compartment in cm$^2$.

Intrinsic currents used in TC and NRT neurons and their conductance values are given in the Tables S3 and S4, respectively. Table S5 lists this information for cortical cells. Passive properties common to neuron models are outlined in the Table S6, while those unique to cortical cells are detailed in Table S7.

*Membrane currents*

Voltage-dependent ion channel currents governing intrinsic membrane potential perturbations were modelled using the Hodgkin-Huxley formalism:

$$I_{int} = \bar{g} m^N h (V_M - E_{int}), \qquad (\boldsymbol{S4})$$

$$\frac{dm}{dt} = \frac{m_\infty - m}{\tau_m}, \qquad (\boldsymbol{S5})$$

$$m_\infty = \frac{\alpha_m}{\alpha_m + \beta_m}, \qquad (\boldsymbol{S6})$$

$$\tau_m = \frac{1}{\alpha_m + \beta_m}, \qquad (\boldsymbol{S7})$$

$$\frac{dh}{dt} = \frac{h_\infty - h}{\tau_h}, \qquad (\boldsymbol{S8})$$

$$h_\infty = \frac{\alpha_h}{\alpha_h + \beta_h}, \qquad (\boldsymbol{S9})$$

$$\tau_h = \frac{1}{\alpha_h + \beta_h}, \qquad (\boldsymbol{S10})$$

where $\bar{g}$ is the maximum conductance in $S/cm^2$, m and h are state variables describing channel activation and inactivation, respectively, $m_\infty$ and $h_\infty$ are the resting state functions describing activation and inactivation, $\tau_m$ and $\tau_h$ are state transition time constants for activation and inactivation, α is the forward rate function in $ms^{-1}$, and β is the backward rate function in $ms^{-1}$. The $Ca^{2+}$- and $Na^{+}$-dependent intrinsic membrane currents followed a similar formalism. These equations apply to intrinsic current descriptions in the Supplementary Appendices A (TC neurons), B (NRT neurons), and C (cortical neurons) unless explicitly stated or replaced by different corresponding equations.

Synaptic currents were described (Appendix D) using a similar formalism with voltage or intracellular ion concentration dependencies replaced by extracellular neurotransmitter concentration dependencies (AMPA, NMDA, $GABA_A$). As for the NMDA channel, the simplification went even further replacing the neurotransmitter concentration by delivering a synaptic event[107]. The NMDA receptor model had neurotransmitter, voltage, and extracellular $Mg^{2+}$ concentration dependencies.

No synaptic plasticity was incorporated into the model. All synapses had the same fixed release probability of 0.8[5,108,109]. Weights associated with each synapse slightly differed among the same type of synapses (standard deviation of 5%) and were pseudo-randomly allocated at the beginning of each simulation. The same applied to synaptic latencies (standard deviation of 20%).

Spontaneous miniature PSP (mPSPs) were generated in all synapses. Their amplitudes were

0.17 mV and 0.083 mV for all cortical AMPA and $GABA_A$ synapses, respectively[110]. mPSPs with 0.5 mV amplitude were used in all thalamic synapses except those of intra-NRT synapses which had an amplitude of 0.25 mV. Both cortical and thalamic mPSPs did not contain NMDA and $GABA_B$ components. mPSPs were generated following an exponential distribution that was dependent on regular synaptic events generated in response to presynaptic action potentials:

$$d(t) = \begin{cases} i_1 n e^{-(t-t_0)}, for\ t - t_0 \leq 1000 \\ i_2 n e^{-(t-t_0)}, for\ t - t_0 > 1000 \end{cases}, \qquad (\boldsymbol{S11})$$

where d is the stimulus delivery delay in ms, t is the time in ms, $t_0$ is the time of the last presynaptic spike in ms, n is the number of the same type synapses on the cell, $i_1$ is the average stimulus delivery delay given a single synapse on a neuron immediately following the presynaptic spike, and $i_2$ is the average stimulus delivery delay given a single synapse on a neuron 1000 ms following the presynaptic spike. For cortical neurons, $i_1$ and $i_2$ were equal to 200/3 and 50/3, respectively, whereas for thalamic cells they were 200 and 100, respectively. A mPSP that was still in the queue of delivery at the time when the new presynaptic spike arrived would have its delivery delay reset.

Intracellular concentration dynamics for $[Ca^{2+}]_i$ and $[Na^+]_i$ were modelled by a simple first-order decay (Destexhe et al., 1993a):

$$\frac{d[Ion]_i}{dt} = -\frac{10000 I_{ion}}{ZFd} + \frac{[Ion]_\infty - [Ion]_i}{\tau_D}, \qquad (\boldsymbol{S12})$$

Where $[Ion]_i$ is the intracellular ion concentration in mM, $I_{ion}$ is the sum of all the transmembrane currents carried by the ion in mA/cm$^2$ (exclude $I_{HVA}$ in TC cells), Z is the valence of the ion, d is the depth of the shell in µm, $[Ion]_\infty$ is the resting intracellular ion concentration in mM, $\tau_D$ is the intracellular ion concentration decay time constant in ms. Decay parameters for $[Ca^{2+}]_i$ and $[Na^+]_I$ are in Table S8.

*EEG signal*

The scalp EEG signal produced by the simulations was estimated based on Bédard et al[111]:

$$V = \frac{1}{4\pi\sigma}\sum_{n=1}^{N}\frac{i_n}{r_n}, \qquad (\boldsymbol{S13})$$

where V is the total sum of N local field potentials (µV) at the surface of the scalp, $\sigma = 0.000355$ mS/µm is the conductivity of the neural tissue[112], $i_n$ is the nth current source in nA, and $r_n$ is the distance from the electrode to the current source in μm. The excitatory current sources associated with neurons in L2/3 were assumed to have a vertical distance of 351.8 μm, 693.4 μm in L4, 1089.4 μm in L5, and 1597.2 μm in L6[82,113]. The inhibitory current sources were assumed to be 500 μm deeper than the excitatory sources to reflect the somato-dendritic spatial distribution differences of the two synapse types. Neighbouring cells in the same cortical layer were equidistant (20 μm) along the horizontal extent of the layer[114]. The EEG calculations were based only on the synapses located on excitatory neurons[114] since they are the main contributors to cortical local field potentials[115].

*Data analyses*

Simulated raw EEG traces were filtered using the Butterworth low-pass filter with 40 Hz and 50 Hz passband and stopband frequencies, respectively. Passband ripple and stopband attenuation parameters were set to 0.5 dB and 10 dB, respectively. EEG auto- and cross-correlations were calculated using Matlab's xcorr function. EEG power was calculated using Matlab's fft function.

Membrane potential histograms were produced using 0.2 mV size bins. During cortical slow (<1 Hz) wave simulations Up- and Down-state durations were estimated for each cell individually with the help of membrane potential histograms. The histogram trough between the two bistability peaks was used as a threshold to roughly split the simulated membrane potential into preliminary periods of Up- and Down-states. Preliminary periods of Up-states longer than 100 ms were deemed to be Up-states and the remaining samples were deemed to be Down-states.

APs were assigned to a particular slow (<1 Hz) oscillation Up-state or a delta/sleep spindle cycle by first low-pass filtering the raw EEG trace using the Butterworth filter. Passband ripple and stopband parameters had the same values as those described earlier. Passband and stopband frequencies were 2 Hz and 3 Hz for slow waves, 5 Hz and 7.5 Hz for delta waves, and 20 and 30 Hz for sleep spindles, respectively. Individual EEG oscillation cycle peaks were then identified. Middle points between two neighbouring cycle peaks were deemed to be cycle borders and APs were assigned to their nearest cycles accordingly.

### Supplementary Appendices

### Appendix A: Intrinsic membrane currents in thalamocortical cell models

This Appendix provides the mathematical descriptions of all intrinsic membrane currents used in TC cell models shown in the equation below:

$$I_{M(TC)} = I_{KL} + I_{NaL} + I_{Na} + I_{K(DR)} + I_T + I_{HVA} + I_h + I_{CAN} + I_{Na(P)} + I_A + I_{K1} + I_{AMPA} + I_{NMDA} + I_{GABAa} + I_{GABAb}. \qquad (\boldsymbol{S14})$$

Their maximum conductances and permeabilities are summarised in Table S3.

The fast transient $Na^+$ current ($I_{Na}$) model was adapted from Traub, Wong, Miles and Michelson (1991)[116]:

$$\alpha_m = \frac{0.32(V_M + 28.9)}{1 - e^{-\frac{V_M+28.9}{4}}}, \qquad (\boldsymbol{S15})$$

$$\beta_m = -\frac{0.28(V_M + 1.9)}{1 - e^{\frac{V_M+1.9}{5}}}, \qquad (\boldsymbol{S16})$$

$$\alpha_h = 0.128e^{-\frac{V_M+25}{18}}, \qquad (\boldsymbol{S17})$$

$$\beta_h = \frac{4}{1 + e^{-\frac{V_M+2}{5}}}, \qquad (\boldsymbol{S18})$$

with N = 3 and $E_{Na}$ = 30 mV. The time constants were temperature dependent with the temperature coefficient $q_{10} = 3^{\frac{T-35}{10}}$, where T is the temperature in degrees of Celsius. Only time constants and not amplitudes were temperature-dependent in thalamic cell models.

The persistent delayed rectifier $K^+$ current ($I_{K(DR)}$) was also adapted from Traub, Wong, Miles and Michelson (1991)[116]:

$$\alpha_m = \frac{0.016(V_M + 2.9)}{1 - e^{-\frac{V_M+2.9}{5}}}, \qquad (\boldsymbol{S19})$$

$$\beta_m = 0.25e^{-\frac{V_M+18}{40}}, \qquad (\boldsymbol{S20})$$

with N = 4 and $E_K$ = -90 mV without the inactivation state h. The temperature coefficient was $q_{10} = 3^{\frac{T-35}{10}}$.

The low voltage activated T-type $Ca^{2+}$ current ($I_T$) was adapted from Williams, Tóth, Turner, Hughes and Crunelli (1997)[117] and described by these equations:

$$I_T = \bar{g}m^2h(V_M - E_{Ca}), \qquad (\boldsymbol{S21})$$

$$m_\infty = \frac{1}{1 + e^{-\frac{V_M+57}{6.2}}}, \qquad (\boldsymbol{S22})$$

$$\tau_m = \begin{cases} e^{\frac{V_M+220.35}{66.6}}, for\ V_M < -57 \\ 2.44 + 0.02506e^{-0.0984(V_M-3)}, for\ V_M \geq -57 \end{cases}, \qquad (\boldsymbol{S23})$$

$$h_\infty = \frac{1}{1 + e^{\frac{V_M+80.5}{6.3}}}, \qquad (\boldsymbol{S24})$$

$$\tau_h = \begin{cases} e^{\frac{V_M+405.8}{66.6}}, for\ V_M < -77 \\ 7.66 + 0.02868e^{-0.1054(V_M-3)}, for\ V_M \geq -77 \end{cases}, \qquad (\boldsymbol{S25})$$

Where $E_{Ca}$ = 180 mV. Both time constants were temperature dependent with $q_{10} = 3^{\frac{T-35}{10}}$. The model behaviour was matched to the experimental voltage-clamp data in Huguenard and Prince(1992)[118].

The non-inactivating HVA $Ca^{2+}$ channels ($I_{HVA}$) were modelled as in McCormick and Huguenard (1992)[119] and Kay and Wong (1987)[120] but were adapted so that they did not activate $I_{CAN}$ as observed in Hughes, Cope, Blethyn and Crunelli (2002)[121]. This fact required a separate $[Ca^{2+}]_i$ pool for $I_{HVA}$. The equations were as follows:

$$I_{HVA} = \bar{P}m^2G(V_M, Ca_o, Ca_i), \qquad (\boldsymbol{S26})$$

$$m_\infty = \frac{1}{1 + e^{-\frac{0.00225F(4.48+V_M)}{R(T-273.15)}}}, \qquad (\boldsymbol{S27})$$

$$\alpha_m = \frac{1.6}{1 + e^{-0.0072(V_M-20)}}, \qquad (\boldsymbol{S28})$$

$$\beta_m = -\frac{0.02(V_M - 7.31)}{1 - e^{\frac{V_M-7.31}{5.36}}}, \qquad (\boldsymbol{S29})$$

$$G(V, Ca_o, Ca_i) = \frac{0.001Z^2F^2V_M\left(Ca_i - Ca_o e^{-\frac{ZFV_M}{R(T+273.15)}}\right)}{1 - e^{-\frac{ZFV_M}{R(T+273.15)}}}, \qquad (\boldsymbol{S30})$$

where $Ca_o$ = 1.5[122-130] and $Ca_i$ are the extracellular and intracellular $Ca^{2+}$ concentrations in mM, respectively, Z = 2 is the valence of calcium ions, F = 96485.309 J is the Faraday constant, R = 8.3144621 J/Kmol is the gas constant, and T is the temperature in degrees of Celsius.. The temperature coefficient was $q_{10} = 3^{\frac{T-21}{10}}$.

$I_h$ was modelled as in Huguenard and McCormick (1992)[131] but was converted into a kinetic scheme to represent $Ca^{2+}$-dependence of the channel as in Destexhe, Bal, McCormick and Sejnowski (1996)[132]. The $Ca^{2+}$-dependence was implemented via $Ca^{2+}$-binding second messenger protein. The whole model is outlined below:

$$I_h = \bar{g}(o_1 + g_{inc}o_2)(V_M - E_h), \qquad (\boldsymbol{S31})$$

$$c \xleftrightarrow{\alpha_m,\beta_m} o_1, \qquad (\boldsymbol{S32})$$

$$p_0 \xleftrightarrow{k_1,k_2} p_1, \qquad (\boldsymbol{S33})$$

$$o_1 \xleftrightarrow{k_3,k_4} o_2, \qquad (\boldsymbol{S34})$$

$$\alpha_m = \frac{m_\infty}{\tau_m}, \qquad (\boldsymbol{S35})$$

$$\beta_m = \frac{(1 - m_\infty)}{\tau_m}, \qquad (\boldsymbol{S36})$$

$$m_\infty = \frac{1}{1 + e^{-\frac{V_M+75}{5.5}}}, \qquad (\boldsymbol{S37})$$

$$\tau_m = 20 + \frac{1000}{e^{\frac{V_M-89.5}{14.2}} + e^{-\frac{V_M+107}{11.6}}}, \qquad (\boldsymbol{S38})$$

$$k_1 = k_2\left(\frac{Ca_{i(inc)}}{Ca_c}\right)^4, \qquad (\boldsymbol{S39})$$

$$k_3 = \frac{k_4 p_1}{p_c}, \qquad (\boldsymbol{S40})$$

where $g_{inc}$ = 0.5 is the $Ca^{2+}$-mediated increase in $G_h$, $E_h$ = -40 mV, c is the proportion of channels in the closed state, $o_1$ is the proportion of channels in the open protein-unbound state, $o_2$ is the proportion of channels in the open protein-bound state, $p_0$ is the proportion of second messenger proteins in the $Ca^{2+}$-unbound state, $p_1$ is the proportion of second messenger proteins in the $Ca^{2+}$-bound state, $k_1$ and $k_2$ = 0.00015 are the $Ca^{2+}$-dependent transition rates between these two protein states in $mM^{-4}ms^{-1}$, $k_3$ and $k_4$ = 0.00007 are the $Ca^{2+}$-bound protein-dependent transition rates between open channel states with different conductances in $ms^{-1}$, $Ca_{i(inc)}$ is the increase in the $[Ca^{2+}]_i$ relative to the resting value in mM, $Ca_c$ = 0.00085 sets the $Ca_{i(inc)}$ threshold value above which $k_1$ functions in the superlinear regime (mM), $p_c$ = 0.017 sets the $p_1$ threshold value above which $k_3$ exceeds $k_4$. $\tau_m$ depended on temperature with $q_{10} = 3^{\frac{T-36}{10}}$. The model behaviour was tested against the experimental voltage clamp data in McCormick and Pape (1990)[133].

The $Ca^{2+}$-activated non-specific cation current ($I_{CAN}$) was implemented using a kinetic scheme with a $Ca^{2+}$-binding second messenger molecule[134-136]:

$$I_{CAN} = \bar{g}m^2h(V_M - E_{CAN}), \qquad (\boldsymbol{S41})$$

$$m_\infty = \frac{0.0001(\frac{Ca_i}{0.00045})^2}{0.0001(\frac{Ca_i}{0.00045})^2 + 0.0001}, \qquad (\boldsymbol{S42})$$

$$h_\infty = \frac{1}{(\frac{Ca_i}{0.00036})^{20} + 1}, \qquad (\boldsymbol{S43})$$

$$\tau_h = \max\left(\left\{\frac{1}{0.00019\left(\frac{0.00036}{Ca_i}\right)^{20} + 0.00019}, 0.1\right\}\right), \qquad (\boldsymbol{S44})$$

where $E_{CAN}$ = 0 mV and $\tau_m$ = 250 ms. $I_{CAN}$ amplitude and dynamics were constrained by the experimental observations in Hughes, Cope, Blethyn and Crunelli (2002)[121].

$I_{Na(P)}$ was modelled according to Parri and Crunelli (1998)[137] but the activation time constant was adopted from the fast transient $Na^+$ channels described by Traub, Wong, Miles and Michelson (1991)[116] but hyperpolarised by 37.68 mV:

$$m_\infty = \frac{1}{1 + e^{-\frac{V_M + 53.87)}{8.57}}}, \qquad (\boldsymbol{S45})$$

$$\alpha_m = \frac{0.32(V_M + 66.58)}{1 - e^{-\frac{V_M + 66.58}{4}}}, \qquad (\boldsymbol{S46})$$

$$\beta_m = -\frac{0.28(V_M + 39.58)}{1 - e^{\frac{V_M + 39.58}{5}}}, \qquad (\boldsymbol{S47})$$

with N = 1, $E_{Na(P)}$ = 30 mV, and h being absent. The temperature coefficient was $q_{10} = 3^{\frac{T-35}{10}}$. $I_{Na(P)}$ amplitude was constrained to be within the experimentally observed range reported by Parri and Crunelli (1998)[137].

$I_A$ model was adopted from Huguenard and McCormick (1992)[131] and was constrained to match the voltage clamp data of Huguenard, Coulter and Prince (1991)[138]:

$$I_A = (\bar{g}_1 m_1^4 h_1 + \bar{g}_2 m_2^4 h_2)(V_M - E_A), \qquad (\boldsymbol{S48})$$

$$m_{1\infty} = \frac{1}{1 + e^{-\frac{V_M + 60}{8.5}}}, \qquad (\boldsymbol{S49})$$

$$m_{2\infty} = \frac{1}{1 + e^{-\frac{V_M + 36}{20}}}, \qquad (\boldsymbol{S50})$$

$$\tau_{m1} = \tau_{m2} = \frac{1}{e^{\frac{V_M+35.8}{19.7}} + e^{-\frac{V_M+79.7}{12.7}}}, \qquad (\boldsymbol{S51})$$

$$h_{1\infty} = h_{2\infty} = \frac{1}{1 + e^{\frac{V_M+78}{6}}}, \qquad (\boldsymbol{S52})$$

$$\tau_{h1} = \begin{cases} \frac{1}{e^{\frac{V_M+46}{5}} + e^{-\frac{V_M+238}{37.5}}}, for\ V_M < -63 \\ 19, for\ V_M \geq -63 \end{cases}, \qquad (\boldsymbol{S53})$$

$$\tau_{h2} = \begin{cases} \tau_{h1}, for\ V_M < -73 \\ 60, for\ V_M \geq -73 \end{cases}, \qquad (\boldsymbol{S54})$$

with $E_A$ = -90 mV and $q_{10} = 3^{\frac{T-23}{10}}$.

$I_{K1}$ model was adapted from Huguenard and Prince (1991)[139]:

$$m_\infty = \frac{1}{1 + e^{-\frac{V_M+5}{8.6}}}, \qquad (\boldsymbol{S55})$$

with N = 1, $E_A$ = -90 mV, $\tau_m$ = 2.5 ms, $q_{10} = 3^{\frac{T-22}{10}}$, and h being absent.

### Appendix B: Intrinsic membrane currents in nucleus reticularis thalami cell models

This Appendix provides mathematical descriptions of all intrinsic membrane currents used in NRT cell models shown in the equation below:

$$I_{M(NRT)} = I_{KL} + I_{NaL} + I_{Na} + I_{K(DR)} + I_{Ts} + I_{HVA} + I_h + I_{AHP} + I_{CAN} + I_{Na(P)} + I_{K[Na]} + I_{AMPA} + I_{NMDA} + I_{GABAa} + I_{gap(1,1)} + I_{gap(1,2)} + I_{gap(2,1)} + I_{gap(2,2)}. \qquad (\mathbf{56})$$

Their maximum conductances and permeabilities are summarised in Table S4.

With a few adjustments most of the intrinsic membrane currents in NRT cells were the same as those used in TC cells. They include $I_{Na}$, $I_{K(DR)}$, $I_{HVA}$, $I_h$, $I_{CAN}$, and $I_{Na(P)}$. $I_{Na}$ voltage dependencies were hyperpolarised relative to TC cells by 8 mV. $I_{K(DR)}$ voltage dependencies were hyperpolarised by 12 mV. With regards to $I_h$, the following parameters were changed: $g_{inc}$ = 2, $k_4$ = 0.00007 $ms^{-1}$, $Ca_c$ = 0.00175 mM, and $p_c$ = 0.017.

The model for the slow T-type $Ca^{2+}$ current ($I_{Ts}$) was described in Huguenard and Prince (1992)[118] with time constants adopted from Destexhe, Contreras, Steriade, Sejnowski and Huguenard (1996)[140]:

$$E_{Ca} = \frac{1000R(T + 273.15)}{2F} log_{10}\left(\frac{Ca_o}{Ca_i}\right), \qquad (\mathbf{S57})$$

$$m_\infty = \frac{1}{1 + e^{-\frac{V_M+50}{7.4}}}, \qquad (\mathbf{S58})$$

$$\tau_m = 3 + \frac{1}{e^{\frac{V_M+25}{10}} + e^{-\frac{V_M+100}{15}}}, \qquad (\mathbf{S59})$$

$$h_\infty = \frac{1}{1 + e^{\frac{V_M+78}{5}}}, \qquad (\mathbf{S60})$$

$$\tau_h = 85 + \frac{1}{e^{\frac{V_M+46}{4}} + e^{-\frac{V_M+405}{50}}}, \qquad (\mathbf{S61})$$

with N = 2, $Ca_o$ = 1.5 mM, and $q_{10} = 3^{\frac{T-24}{10}}$.

The $I_{AHP}$ model was outlined in Xia, Fakler, Rivard, Wayman, Johnson-Pais, Keen, Ishii, Hirschberg, Bond, Lutsenko, Maylie and Adelman (1998)[141] and calibrated by data in Cueni, Canepari, Lujan, Emmenegger, Watanabe, Bond, Franken, Adelman and Luthi (2008)[142]:

$$I_{AHP} = (\bar{g}_1 m_{1\infty} + \bar{g}_2 m_{2\infty})(V_M - E_{CAN}), \qquad (\mathbf{S62})$$

$$m_{1\infty} = \frac{1}{\left(\frac{Ca_{EC50,1}}{Ca_{i(inc)}}\right)^{5.3} + 1}, \qquad (\boldsymbol{S63})$$

$$m_{2\infty} = \frac{1}{\left(\frac{Ca_{EC50,2}}{Ca_{i(inc)}}\right)^{5.3} + 1}, \qquad (\boldsymbol{S64})$$

with $E_A$ = -90 mV, $Ca_{EC50,\,1}$ = 0.001, and $Ca_{EC50,\,2}$ = 0.00032 are $[Ca^{2+}]_i$ of the half-maximal response (mM) for the two components, $\tau_{m1}$ = 15 ms, $\tau_{m2}$ = 830 ms, $q_{10} = 3^{\frac{T-34.25}{10}}$, and h being absent.

The $Ca^{2+}$-activated non-specific cation current ($I_{CAN}$) was implemented following the scheme outlined for the TC cells but with a few differences. Modified equations are shown below:

$$I_{CAN} = \bar{g}m^2(V_M - E_{CAN}), \qquad (\boldsymbol{S65})$$

$$m_\infty = \frac{0.000004(\frac{Ca_i}{0.00045})^2}{0.000004(\frac{Ca_i}{0.00045})^2 + 0.000004}, \qquad (\boldsymbol{S66})$$

$$\tau_m = \frac{1}{0.000004(\frac{Ca_i}{0.00045})^2 + 0.000004}, \qquad (\boldsymbol{S67})$$

where $E_{CAN}$ = 0 mV.

$I_{K[Na]}$ was taken from Bischoff, Vogel and Safronov (1998)[143]:

$$m = \frac{1}{1 + (\frac{38.7}{Na_i})^{3.5}}, \qquad (\boldsymbol{S68})$$

With N = 1, $E_{K[Na]}$ = -90 mV, $q_{10} = 2.3^{\frac{T-37}{10}}$ for the amplitude, and h being absent.

### Appendix C: Intrinsic membrane currents in neocortical cell models

This Appendix provides mathematical descriptions of all intrinsic membrane currents used in the cortical cell models shown in the equations below:

$$I_S = I_{KL} + I_{NaL} + I_{Na} + I_{K(DR)} + I_{Na(P)} + I_{K[Na]} + I_{GABAa} + I_{GABAb} + I_{DS}, \qquad (\boldsymbol{S69})$$

$$I_D = I_{KL} + I_{NaL} + I_{Na} + I_A + I_M + I_{fAHP} + I_{sAHP} + I_h + I_{Na(P)} + I_{K[Na]} + I_T + I_{HVA} + I_{AMPA} + I_{NMDA} + I_{SD}, \qquad (\boldsymbol{S70})$$

Most of the models of cortical currents were used previously in Mainen and Sejnowski (1996)[144]. Their maximum conductances and permeabilities are summarised in Table S5.

$I_{Na}$ is included in both axosomatic and dendritic compartments and was originally taken from Mainen and Sejnowski (1996)[144]:

$$\alpha_m = \frac{0.182(V_M + 25)}{1 - e^{-\frac{V_M+25}{9}}}, \qquad (\boldsymbol{S71})$$

$$\beta_m = -\frac{0.124(V_M + 25)}{1 - e^{\frac{V_M+25}{9}}}, \qquad (\boldsymbol{S72})$$

$$\alpha_h = \frac{0.024(V_M + 40)}{1 - e^{-\frac{V_M+40}{5}}}, \qquad (\boldsymbol{S73})$$

$$\beta_h = -\frac{0.0091(V_M + 65)}{1 - e^{-\frac{V_M+65}{5}}}, \qquad (\boldsymbol{S74})$$

$$\tau_h = \frac{1}{1 + e^{\frac{V_M+55}{6.2}}}, \qquad (\boldsymbol{S75})$$

with N = 3 and $E_{Na}$ = 60 mV.

$I_{K(DR)}$ model was used only in the axosomatic compartment and was adopted from the same source:

$$\alpha_m = \frac{0.02(V_M - 25)}{1 - e^{-\frac{V_M-25}{9}}}, \qquad (\boldsymbol{S76})$$

$$\beta_m = -\frac{0.002(V_M - 25)}{1 - e^{\frac{V_M-25}{9}}}, \qquad (\boldsymbol{S77})$$

with N = 1 and h being absent. Amplitudes and time constants of both $I_{Na}$ and $I_{K(DR)}$ increased and decreased with temperature, respectively. The temperature factor was $q_{10} = 2.3^{\frac{T-23}{10}}$.

$I_{Na(P)}$ was expressed in both compartments and adopted from Mainen and Sejnowski (19960[144] with the time constant taken from Timofeev, Grenier, Bazhenov, Sejnowski and Steriade (2000)[145]:

$$m_{\infty} = \frac{1}{1 + e^{-\frac{V_M + 42}{5}}}, \qquad (\boldsymbol{S78})$$

with N = 1, $E_{Na(P)}$ = 60 mV, $\tau_m$ = 0.05 ms, $q_{10} = 2.3^{\frac{T-36}{10}}$ for the amplitude and the time constant, and h being absent.

$I_{K[Na]}$ was also expressed in both compartments and was taken from Bischoff, Vogel and Safronov (1998)[143]:

$$m = \frac{1}{1 + (\frac{38.7}{Na_i})^{3.5}}, \qquad (\boldsymbol{S79})$$

with N = 1, $E_{K[Na]}$ = -90 mV $q_{10} = 2.3^{\frac{T-37}{10}}$ for the amplitude, and h being absent.

$I_A$ was expressed in the dendritic compartment, modelled according to Keren, Peled and Korngreen (2005)[146] and constrained against the experimental data of Korngreen and Sakmann (2000)[147]:

$$m_{\infty} = \frac{1}{1 + e^{-\frac{V_M + 47}{29}}}, \qquad (\boldsymbol{S80})$$

$$\tau_m = 0.34 + 0.92e^{-(\frac{V_M + 71}{59})^2}, \qquad (\boldsymbol{S81})$$

$$h_{\infty} = \frac{1}{1 + e^{\frac{V_M + 66}{10}}}, \qquad (\boldsymbol{S82})$$

$$\tau_h = 8 + 49e^{-(\frac{V_M + 73}{23})^2}, \qquad (\boldsymbol{S83})$$

with N = 4, $E_A$ = -90 mV, and $q_{10} = 2.3^{\frac{T-21}{10}}$ for the amplitude and time constants.

Similarly, $I_M$ was localised within the dendritic compartment and modelled according to Mainen and Sejnowski (1996)[144] and Yamada, Koch and Adams (1989)[148]:

$$\alpha_m = \frac{0.0001(V_M + 30)}{1 - e^{-\frac{V_M + 30}{9}}}, \qquad (\boldsymbol{S84})$$

$$\beta_m = -\frac{0.0001(V_M + 30)}{1 - e^{\frac{V_M + 30}{9}}}, \qquad (\boldsymbol{S85})$$

with N = 1, $E_A$ = -90 mV, $q_{10} = 2.3^{\frac{T-23}{10}}$ for the amplitude and the time constant, and h being absent.

$I_{fAHP}$ was expressed in the dendritic compartment only and adopted from Mainen and Sejnowski (1996)[144] with α = 10$Ca_i$, β = 0.02, with N = 1, $E_A$ = -90 mV, $q_{10} = 2.3^{\frac{T-23}{10}}$ for the amplitude and the time constant, and h being absent. Meanwhile $I_{sAHP}$ was based on a model derived in the context of non-cortical cells[141,142] with equations being the same as in NRT cells (see Equations S62-64). Changes were: $\bar{g}_2$ = 0.000001-0.00145 S/cm$^2$. $I_{sAHP}$ was present only in dendritic compartments of ND cells.

$I_h$ was taken from Keren, Peled and Korngreen (2005)[146] with the $Ca^{2+}$-dependence modelled similarly to thalamic cells[132]. The equations were also the same except for $m_\infty$, $\tau_m$, and $k_1$ which were:

$$m_\infty = \frac{1}{1+e^{-\frac{V_M+91}{6}}}, \qquad (\boldsymbol{S86})$$

$$\tau_m = \frac{1}{0.0004e^{-0.025V_M}+0.088e^{0.062V_M}}, \qquad (\boldsymbol{S87})$$

$$k_1 = k_2\left(\frac{Ca_i}{Ca_c}\right)^4. \qquad (\boldsymbol{S88})$$

Other parameters were $E_{K[Na]}$ = -30 mV, $k_2$ = 0.00015 mM$^{-4}$ms$^{-1}$, $k_4$ = 0.00007 ms$^{-1}$, $Ca_c$ = 0.0015 mM, $p_c$ = 0.017, $q_{10} = 3.5^{\frac{T-36}{10}}$ for the time constant only, and h being absent.

$I_T$ was based on Destexhe, Neubig, Ulrich and Huguenardv (1998)[149] and described by Goldman-Hodgkin-Katz equations:

$$I_T = \bar{P}m^2hG(V_M, Ca_o, Ca_i), \qquad (\boldsymbol{S89})$$

$$m_\infty = \frac{1}{1+e^{-\frac{V_M+57}{6.2}}}, \qquad (\boldsymbol{S90})$$

$$\tau_m = 0.612 + \frac{1}{e^{-\frac{V_M+132}{16.7}}+e^{\frac{V_M+16.8}{18.2}}}, \qquad (\boldsymbol{S91})$$

$$h_\infty = \frac{1}{1+e^{\frac{V_M+81}{4}}}, \qquad (\boldsymbol{S92})$$

$$\tau_h = \begin{cases} e^{\frac{V_M+467}{66.6}}, for\ V_M \le -80 \\ 28 + e^{-\frac{V_M+22}{10.5}}, for\ V_M > -80 \end{cases}, \qquad (\boldsymbol{S93})$$

where $\overline{\mathrm{P}}$ is the maximum membrane permeability to $Ca^{2+}$ in cm/s ($\overline{\mathrm{P}}_{RS,EF}$ = 0.000001 and $\overline{\mathrm{P}}_{IB,RIB,ND}$ = 0.00001). Both time constants were temperature dependent with $q_{10} = 3^{\frac{T-24}{10}}$. $I_T$ was absent in FS cells.

$I_{HVA}$ was modelled according to Mainen and Sejnowski (1996)[144]:

$$\alpha_m = \frac{0.055(V_M + 27)}{1 - e^{-\frac{V_M+27}{3.8}}}, \qquad (\boldsymbol{S94})$$

$$\beta_m = 0.94e^{-\frac{V_M+75}{17}}, \qquad (\boldsymbol{S95})$$

$$\alpha_h = 0.000457e^{-\frac{V_M+13}{50}}, \qquad (\boldsymbol{S96})$$

$$\beta_h = \frac{0.0065}{1 + e^{-\frac{V_M+15}{28}}}, \qquad (\boldsymbol{S97})$$

with N = 2, $E_{HVA}$ = 140 mV, $Ca_o$ = 1.5 mM, and $q_{10} = 2.3^{\frac{T-23}{10}}$ for the amplitude and time.

### Appendix D: Synaptic membrane current models

This appendix provides the mathematical descriptions of synaptic current models and their parameters used in the corticothalamic network model.

Except the NMDA component, AMPA, $GABA_A$, and $GABA_B$ postsynaptic currents were modelled based on a simplifying assumption of the neurotransmitter concentration dynamics in the synaptic cleft as a unitary amplitude pulse as described in Destexhe, Mainen and Sejnowski (1994)[150]:

$$I_{AMPA/GABA_A} = \bar{g}m\left(V_M - E_{AMPA/GABA_A}\right), \qquad (\boldsymbol{S98})$$

$$m = \begin{cases} m_\infty + (m(t_0) - m_\infty)e^{-\frac{t-t_0}{\tau_m}}, for\ t - t_0 \leq T_{dur} \\ m(t_0 + T_{dur})e^{-\beta(t-t_0-T_{dur})}, for\ t - t_0 > T_{dur} \end{cases}, \qquad (\boldsymbol{S98})$$

$$m_\infty = \frac{\alpha T_{max}}{\alpha T_{max} + \beta}, \qquad (\boldsymbol{S100})$$

$$\tau_m = \frac{1}{\alpha T_{max} + \beta}, \qquad (\boldsymbol{S101})$$

where $I_{AMPA/GABAa}$ is the postsynaptic current in nA, $\bar{g}$ is the maximal conductance in µS, $t_0$ is the onset time of the neurotransmitter pulse (ms), $T_{dur}$ is the duration of the neurotransmitter pulse (ms), $T_{max}$ is the amplitude of the pulse in mM. Tables S10 and S11 summarise AMPAR and $GABA_AR$ parameter sets used in this model.

Thalamic and cortical postsynaptic $GABA_B$ currents were based on the same simplifying solution but involving a second messenger protein as outlined in Destexhe, Bal, McCormick and Sejnowski (1996)[132] and Thomson and Destexhe (1990)[40]:

$$\frac{dR}{dt} = k_1 T_{max}(1 - R) - k_2 R, \qquad (\boldsymbol{S102})$$

$$\frac{dP}{dt} = k_3 R - k_4 P, \qquad (\boldsymbol{S103})$$

$$I_{GABA_B} = \bar{g}m\frac{P^4}{P^4 + K_d}\left(V_M - E_{GABA_B}\right), \qquad (\boldsymbol{S104})$$

where $E_{GABAb}$ = -90 mV, R is the fraction of activated receptor, P is the concentration of activated second messenger protein in mM, $K_d$ is the dissociation constant of the binding of the activated protein on the $K^+$ channels mediating the $GABA_B$ postsynaptic current in $mM^4$, $k_1$ ($mM^{-1}ms^{-1}$) and $k_2$ ($ms^{-1}$) are the forward and backward receptor state transition rates, respectively, whereas $k_3$ ($ms^{-1}$) and $k_4$ ($ms^{-1}$) are second messenger protein activation and inactivation rates, respectively. The transmitter $T_{max}$ is only present for a limited period $T_{dur}$. The sets of $GABA_BR$ parameters are outlined in Table S12.

The NMDA postsynaptic current model in the thalamus and the cortex was the most complex of all other synaptic channels used here and was based on the work presented in Moradi, Moradi, Ganjkhani, Hajihasani, Gharibzadeh and Kaka (2013)[107] but excluding short-term depression. The following is the outline:

$$I_{NMDA} = \bar{g}(f_{VI} + f_{VD})(w_C C + w_B B - A)Mg(V_M - E_{NMDA}), \qquad (\boldsymbol{S105})$$

$$\frac{\partial f_{VD}}{\partial t} = -\frac{f_{VI}(w_C C + w_B B)(f_{VD,\infty} - f_{VD})}{\tau_g}, \qquad (\boldsymbol{S106})$$

$$\tau_g = \frac{7}{q_{10,g}}, \qquad (\boldsymbol{S107})$$

$$q_{10,g} = 1.52^{\frac{(T-26)}{10}}, \qquad (\boldsymbol{S108})$$

$$g_{VD,\infty} = k(V_M - V_0), \qquad (\boldsymbol{S109})$$

$$\frac{dA}{dt} = -\frac{A}{\tau_A}, \qquad (\boldsymbol{S110})$$

$$\tau_A = \frac{\tau_{A,0} + a_A e^{-\lambda_A V_M}}{q_{10,A}}, \qquad (\boldsymbol{S111})$$

$$q_{10,A} = 2.2^{\frac{(T-35)}{10}}, \qquad (\boldsymbol{S112})$$

$$\frac{dB}{dt} = -\frac{B}{\tau_B}, \qquad (\boldsymbol{S113})$$

$$\tau_B = \frac{\tau_{B,0} + a_B(1 - e^{-\lambda_B V_M})}{q_{10,B}}, \qquad (\boldsymbol{S114})$$

$$q_{10,B} = 3.68^{\frac{(T-35)}{10}}, \qquad (\boldsymbol{S115})$$

$$\frac{dC}{dt} = -\frac{C}{\tau_C}, \qquad (\boldsymbol{S116})$$

$$\tau_C = \frac{\tau_{C,0} + a_C(1 - e^{-\lambda_C V_M})}{q_{10,C}}, \qquad (\boldsymbol{S117})$$

$$q_{10,C} = 2.65^{\frac{(T-35)}{10}}, \qquad (\boldsymbol{S118})$$

$$Mg = \frac{1}{1 + \frac{Mg_o}{Mg_{IC50}} e^{-\frac{0.001 Z \delta F V_M}{R(T+273.15)}}}, \qquad (\boldsymbol{S119})$$

where $E_{NMDA}$ = -0.7 mV, $f_{VI}$ = 0.5 and $f_{VD}$ are the voltage-independent and voltage-dependent channel conductance components in fractions, respectively, $f_{VD,\infty}$ is the resting voltage-dependent conductance function (µS), $V_0$ = -100 mV is the baseline $V_M$ at which the $f_{VD}$ = 0, k =0.007 $mV^{-1}$ is the factor relating $V_M$ change to the $f_{VD}$, $\tau_g$ is the voltage-dependent conductance transition time constant (ms), A is the channel activation state dependent on the neurotransmitter, B and C are the deactivation states, $w_B$ = 0.65 and $w_C$ = 0.35 set the proportions of the two inactivation terms ($w_B + w_C = 1$), Mg determines the $Mg^{2+}$ block, $\tau_A$, $\tau_B$, and $\tau_C$ are the activation and the two deactivation time constants (ms), $\tau_{A,0}$ = 3 ms, $\tau_{B,0}$ = 25.057 ms, and $\tau_{C,0}$ = 232.27 ms are initial time constants (ms) at $V_m$ = 0 mV, $a_A$ = 1, $a_B$ = 2.2364, and $a_C$ = 43.495 are tuning factors, $\lambda_A$ = 1, $\lambda_B$ = 0.0243, and $\lambda_C$ = 0.01 are decay constants, $Mg_o$ is the extracellular $Mg^{2+}$ concentration in mM, $Mg_{IC50}$ = 4.1 mM is the 50% $Mg^{2+}$ inhibition concentration in mM at $V_m$ = 0 mV, Z = 2 is the valence of $Mg^{2+}$, and $\delta$ = 0.8 is the relative electrical distance of the binding site of $Mg^{2+}$ from the outside of the membrane. The sets of NMDAR parameters are outlined in Table S13.
